# A Neurocompliant Ionic Conductive Hydrogel Orchestrates Glial and Vascular Remodeling to Mitigate Traumatic Brain Injury

**DOI:** 10.64898/2026.10.29.762869

**Authors:** Prince D. Okoro, Razieh Jaberi, Shaylyn Blackburn, Dhyan Patel, Nadeem Halasah, Jacob Mapa, Michael Shih, Ryan Botzbach, Jinrui Tan, Valentine I. Vullev, Huinan Liu, Vijayalakshmi Santhakumar, Iman Noshadi

## Abstract

Developing conductive scaffolds for neural regeneration is often hindered by the cytotoxicity and mechanical mismatch inherent to conventional electronic fillers. Here, we report an ultrasoft ionically conductive hydrogel engineered by grafting choline-based bioionic liquids (BIL) onto fish gelatin (FiBGel). Systematic optimization yields brain-mimetic conductivity (0.25–0.68 S m⁻¹), elastic moduli (0.6–0.9 kPa), and physiological water content (72–81%). In 3D culture, the matrix drives spontaneous neurogenesis, achieving ∼90% MAP2+ neurons with robust calcium signaling, while concurrently supporting endothelial network formation. We report the first intracerebral administration of a BIL hydrogel, demonstrating remarkable biocompatibility and immunomodulation. In a traumatic brain injury model, a single FiBGel dose halves reactive astrogliosis, enhances cortical revascularization, and limits lesion cavity expansion. Leveraging an FDA-recognized micronutrient, FiBGel offers a translatable strategy that integrates neuromechanics, conductivity, immune modulation, and vascular support within one injectable scaffold for minimally invasive treatment of brain injuries.

## 1. Introduction

Brain tissue repair and regeneration underpin outcomes across the treatments of numerous neurological disorders, including and traumatic brain injury (TBI), stroke, and neurodegenerative diseases, where the failure of endogenous repair mechanisms results in permanent dysfunction and disability. TBI represents among the foremost causes of death and disability, burdening roughly 1.5 million Americans and 69 million global cases annually[1]. Unlike the peripheral nervous system, where Schwann cells rapidly adopt a repair phenotype to coordinate axonal regrowth and remyelination, the adult central nervous system (CNS) possesses limited endogenous regenerative potential due to the absence of pro-regenerative glial programs and rapid formation of a glial scar that inhibits axon extension[2,3]. Despite improved acute care, deficits persist as astrogliosis propagates beyond lesions, reinforcing barriers to circuit reconstitution[4,5]. Consequently, exogenous brain-like matrices that temper neuroinflammation while preserving or restoring neural signaling pathways are compelling adjuncts to current therapies.

Neural tissue engineering offers promising routes to design microenvironments that recapitulate brain-specific cues influencing cell fate and tissue development, particularly endogenous electrical fields that regulate neural differentiation, axon guidance, and circuit assembly. Conventional conductive hydrogels rely on (1) electron- and hole-conductivity pathways along polypyrrole (PPy)[6], polyaniline (PANI)[7], or carbon nanomaterials[8,9], or (2) ionic conductivity mediated by poly(3,4-ethylenedioxythiophene):polystyrene sulfonate (PEDOT:PSS)[10], which ensures efficient charge transport but raises significant biosafety and biodegradation concerns. Notably, graphene and carbon nanotubes demonstrate cytotoxicity and genotoxicity at concentrations as low as 0.002–0.05% (w/v)[11,12], while neither PEDOT nor PSS contains biodegradable linkages[13], precluding scaffold resorption and compounding chronic foreign-body risks. Moreover, these additives commonly stiffen matrices, worsening the mechanical mismatch with native brain tissue that promotes gliosis *in vivo*[14,15], underscoring the need for safe neurocompliant conductive alternatives.

Ionically conductive hydrogels represent a paradigm shift, mediating charge transport where mobile ions are the charge carriers in aqueous-polymer networks. It fundamentally aligns with the electrochemical basis of neural tissues. Unlike electronic conductors, these materials mirror native neural signaling where ionic fluxes mediate extracellular communication[16,17], involving conformal low-impedance contacts while mitigating faradaic side reactions. Bio-ionic liquids (BILs) exemplify this approach, comprising biocompatible ionic media formed by strategic pairing of organic cations with biogenic anions to achieve systematically tunable electrical and physicochemical properties[18]. Choline-based BILs are particularly attractive ion carriers as choline serves as an essential micronutrient for phospholipid synthesis[19], acetylcholine production[20], and neurodevelopment[21]. Moreover, cholinium groups can engage α7 nicotinic acetylcholine receptors to activate the cholinergic anti-inflammatory pathway[22–24] and endothelial pro-angiogenic signaling[25–27], providing mechanistic routes for both immune modulation and vascular support.

We previously established BIL-conjugated porcine gelatin methacryloyl (BioGel) hydrogels as a versatile platform with tunable conductivity, mechanics, hydration, and degradation while maintaining excellent biocompatibility *in vitro* and *in vivo*[28–30]. These features allow attaining multifunctional hydrogels with adhesive, hemostatic, antimicrobial, and angiogenic properties[31,32], recently demonstrating therapeutic efficacy in chronic wound healing[33]. Related BIL systems show neurocompatibility *in vitro*[34,35], yet most target peripheral nerve repair with elastic moduli (5–17 kPa) exceeding the 0.1–2 kPa range of brain tissue[36]. Such mechanical mismatch drives chronic microglial/astrocytic responses and blood–brain barrier disruption[14,15]. Achieving brain-mimetic stiffness must be coupled with physiologically relevant conductivity while preserving high water content that sustains ionic homeostasis and metabolite transport within the peri-implant milieu. For successful deep-brain applications, BIL hydrogels must simultaneously satisfy these integrated property requirements to ensure optimal tissue integration and therapeutic efficacy.

Here we design a neurocompliant ionically conductive hydrogel by covalently grafting choline BIL onto an ultrasoft fish gelatin methacryloyl (GelMA) backbone. By systematically varying the ionic loading, we tune mechanical strength, ionic conductivity, water content, and degradation rates toward the mechanical and electrochemical milieu of the cortical gray matter. We interrogate how this physicochemical landscape affects human neural progenitor viability, lineage commitment, and endothelial tubulogenesis *in vitro*, and probe the capacity of the material to coordinate neurogenic and angiogenic processes. This work reports the first intracerebral administration of a BIL hydrogel and establishes acute biocompatibility. As a proof of concept, we validate the therapeutic potential of the optimized formulation in a moderate traumatic brain injury model, utilizing neuroimmune activation and microvascular formation as key metrics to establish its utility as an injectable brain-matched scaffold for minimally invasive neuroregenerative therapy.

## 2. Results

### 2.1. Design rationale, synthesis, and characterization of bioionic hydrogels

To bridge the disparity between electronic conductors and native biological signaling, we engineered a neurocompliant hydrogel interface by integrating choline-based bioionic liquids into a fish GelMA network. We have previously demonstrated that choline-based BILs serve as conductive, adhesive, and bioactive substrates for tissue regeneration when conjugated with porcine-based GelMA [28,29,31–33]. However, resorting to softer matrices than previously reported is key for matching the stiffness of native brain tissue (0.1-2 kPa)[36–38], with substrates ≤1 kPa specifically enhancing neural stem cell growth and differentiation [39–41]. Unconfined compression testing of 4%, 7%, and 10% (w/v) fish GelMA versus porcine GelMA reveals distinct stress-strain curves and consistently lower compressive moduli for fish GelMA (Figure S1). Notably, at the 7% (w/v) concentration, fish GelMA exhibited a compressive modulus of 0.90 ± 0.02 kPa compared to 5.91 ± 0.43 kPa for porcine GelMA, placing the former precisely within the neurocompliant range. Rheological profiling further confirms that fish GelMA resists thermoresponsive gelation from 40 to 0 °C (Figure S2), maintaining a stable liquid state that streamlines cell encapsulation, bioprinting, and minimally invasive delivery.

Choline bitartrate (ChBT) was functionalized with photo-crosslinkable acrylate groups via acid-catalyzed esterification to integrate BIL moieties in the polymer network as previously reported[29,33,42]. ^1^H NMR spectroscopy confirmed the synthesis of acrylated choline bitartrate, evidencing acrylate vinyl protons resonating at δ 6.4 and 5.5 ppm (Figure S3B). The characteristic cholinium resonances persisted after acrylation, with multiplets centered at δ 3.9 ppm (O–CH_2_, 2H) and δ 3.42 ppm (N^+^–CH_2_, 2H) arising from vicinal coupling between the adjacent methylene groups, together with a prominent singlet at δ 3.23 ppm (N^+^(CH_3_)_3_, 9H). Notably, the bitartrate methine doublet at δ 4.4 ppm remained detectable after ethyl acetate washing. Since unreacted tartaric acid is removed during purification, this retention confirms the preservation of the counter-ion required to maintain charge neutrality of the acryloyl-choline cation. Methacrylation of the fish-gelatin backbone was evidenced by new vinyl doublets at δ 5.63 and 5.39 ppm together with a sharp α-methyl singlet at δ 1.9 ppm(Figure S3B), verifying grafting of methacryloyl groups onto ε-lysine and aligning with commonly reported GelMA spectra[43,44].

A facile one-pot strategy was employed for bioionic liquid-conjugated fish gelatin methacryloyl (FiBGel) hydrogel fabrication. It involves combining 7% (w/v) fish GelMA, acrylated ChBT at defined concentrations (2, 4, or 6% w/v), and the photoinitiator LAP (lithium phenyl-2,4,6-trimethylbenzoylphosphinate; 0.25% w/v) (Figure 1A). Precursor solutions were adjusted to physiological pH, and crosslinking was achieved via 405 nm light irradiation for 30 s using LAP photoinitiator (Figure S3A). This approach enables simultaneous network formation and covalent incorporation of bioionic moieties through visible light activation. Visual observation confirmed sol-gel transition, and ^1^H NMR analysis verified BIL incorporation through persistent cholinium resonances at δ 3.9, 3.4, and 3.23 ppm, alongside the unique bitartrate peak at δ 4.4 ppm (Figure S3B). Morphologically, scanning electron microscopy (SEM) reveals that FiBGel maintains an open, highly porous network architecture essential for nutrient transport and neurite infiltration (Figure 1B). For brevity, fish GelMA and acrylated ChBT are hereafter referred to as GelMA and ChBT, respectively.

**Figure 1.**
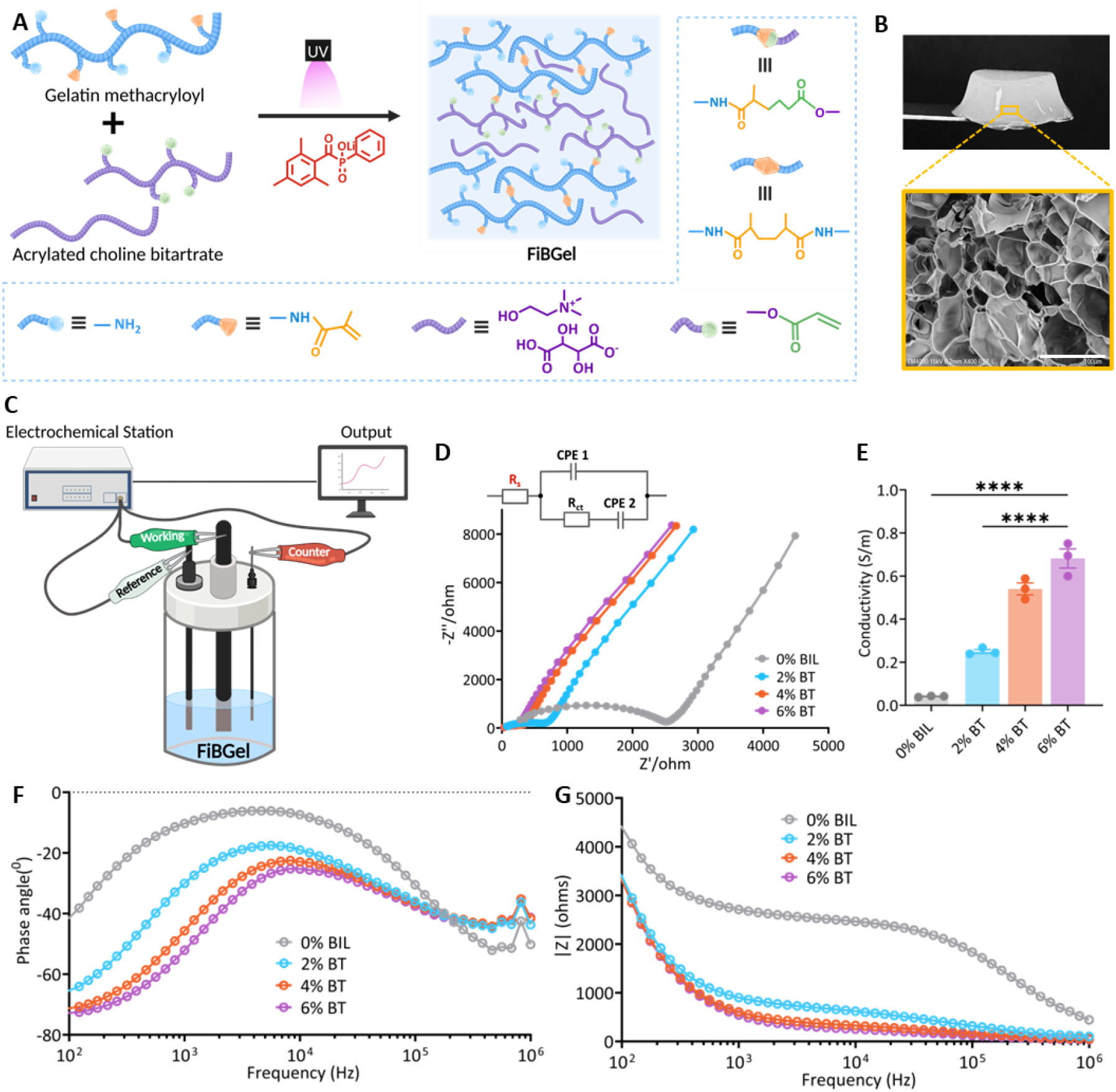
FiBGel synthesis and electrochemical characterization. (A) Fish gelatin methacryloyl (GelMA) and acrylated choline bitartrate (ChBT) were crosslinked via lithium phenyl-2,4,6-trimethylbenzoylphosphinate (LAP) upon irradiation at 405 nm to form bioionic liquid (BIL)-conjugated fish GelMA (FiBGel). (B) Macroscopic images of the crosslinked hydrogels with corresponding scanning electron microscopy (SEM) images highlighting their porous microstructures. Scale bars: 100 μm. (C) Schema of the three-electrode setup used for electrochemical impedance spectroscopy (EIS) and conductivity measurements. (D) Nyquist plots for GelMA hydrogels functionalized with 0, 2, 4, or 6% (w/v) ChBT. Inset shows the modified Randles model with constant phase elements, R(Q(RQ)), used to fit spectra and extract Rs while capturing interfacial polarization. (E) Ionic conductivity calculated from impedance data, demonstrating a significant, concentration-dependent increase with BIL content. (F) Bode plots showing phase angle versus frequency for FiBGel hydrogels, detailing their frequency-dependent electrochemical response across all BIL concentrations. (G) Impedance magnitude |Z| versus frequency for FiBGel formulations, confirming reduced impedance with increasing BIL content. All hydrogel formulations contained 7% (w/v) GelMA and were photocrosslinked using 0.25% (w/v) LAP. The 0% (w/v) BIL samples contain only GelMA and serve as the control. Values represent mean ± SEM. Statistical significance via one-way ANOVA with Tukey’s post-hoc correction (*p ≤ 0.05, **p ≤ 0.01, ***p ≤ 0.001, ****p ≤ 0.0001).

### 2.2. Electrochemical Impedance Profiling and Ionic Conductivity

Regenerating damaged neural tissue demands biomaterials that mimic the native brain’s electrical environment, providing low-resistance pathways for cell-to-cell electrochemical signaling that guide axonal extension and network maturation. To quantify the frequency-dependent electrochemical behavior of the hydrogels, we performed electrochemical impedance spectroscopy (EIS) on *in situ* photocrosslinked hydrogels (Figure 1C & S4A). The Nyquist plots display a systematic leftward shift and reduction in arc diameter with increasing bioionic liquid concentration, indicating a simultaneous decline in bulk resistance (R_S_) and charge transfer resistance (R_CT_) (Figure 1D). We modelled these spectra using a modified Randles circuit (More details in Supplementary Note 1), which incorporates a constant-phase element (CPE) to resolve the non-ideal capacitive behavior arising from the porous hydrogel-electrode interface[45].

Ionic conductivity was calculated from the R_S_ extracted via circuit fitting (Figure S4C), using a cell constant (K) determined via calibration with KCl standards of known conductivity (Figure S4B). The conductivity values revealed a pronounced dependence on choline bitartrate loading (Figure 1E). While the pristine GelMA control displayed a baseline conductivity of 0.041 ± 0.003 S m⁻¹, the incorporation of 2% (w/v) acrylated ChBT elevated this to 0.25 ± 0.02 S m⁻¹, and rose to 0.540 ± 0.048 S m⁻¹ and 0.681 ± 0.076 S m⁻¹ for the 4% and 6% formulations, respectively. This represents a 16-fold enhancement over the unfunctionalized control. Importantly, this tunable range remains well below the conductivity of cerebrospinal fluid (∼1.7–1.8 S m⁻¹)[46], ensuring the material facilitates tissue-level ionic coupling without short-circuiting local field potentials. Bode phase plots (Figure 1F) corroborate this trend, showing the principal dispersion shifting to higher frequencies with increasing ChBT content, which signifies shortened relaxation times and faster ion mobility. Furthermore, the total impedance magnitude (|Z|) decreased systematically with both frequency and ionic liquid concentration (Figure 1G), confirming the establishment of low-resistance pathways essential for effective neural coupling.

### 2.3. Mechanical Properties and Hydrolytic Stability

Viscoelastic profiling is essential as hydrogel stiffness and damping steer neural lineage specification and mitigate foreign-body responses; mechanically matched hydrogels have recently been reported to reduce astrocytic scar thickness by up to 60% [14]. Frequency-sweep rheology (0.1–10 Hz) confirmed that all FiBGel formulations behave as stable, elastic networks with storage moduli (G’) consistently dominating over loss moduli (G’’) and no crossover points across the frequency range (Figure 2A). Incorporating ChBT induced a monotonic, concentration-dependent increase in stiffness relative to GelMA controls. Specifically, the shear modulus (G’) at 1 Hz rose marginally from 0.21 ± 0.01 kPa (2% w/v) to 0.29 ± 0.03 kPa (6% w/v) (Figure 2A). Applying the incompressibility relation for soft tissues (E ∼ 3G’), these values correspond to elastic moduli of 0.63–0.87 kPa (Figure 2B). Despite this stiffening, the damping factor (tan δ) remained ≤ 0.2 for all formulations, confirming effective elastic shape retention (Figure 2C).

**Figure 2.**
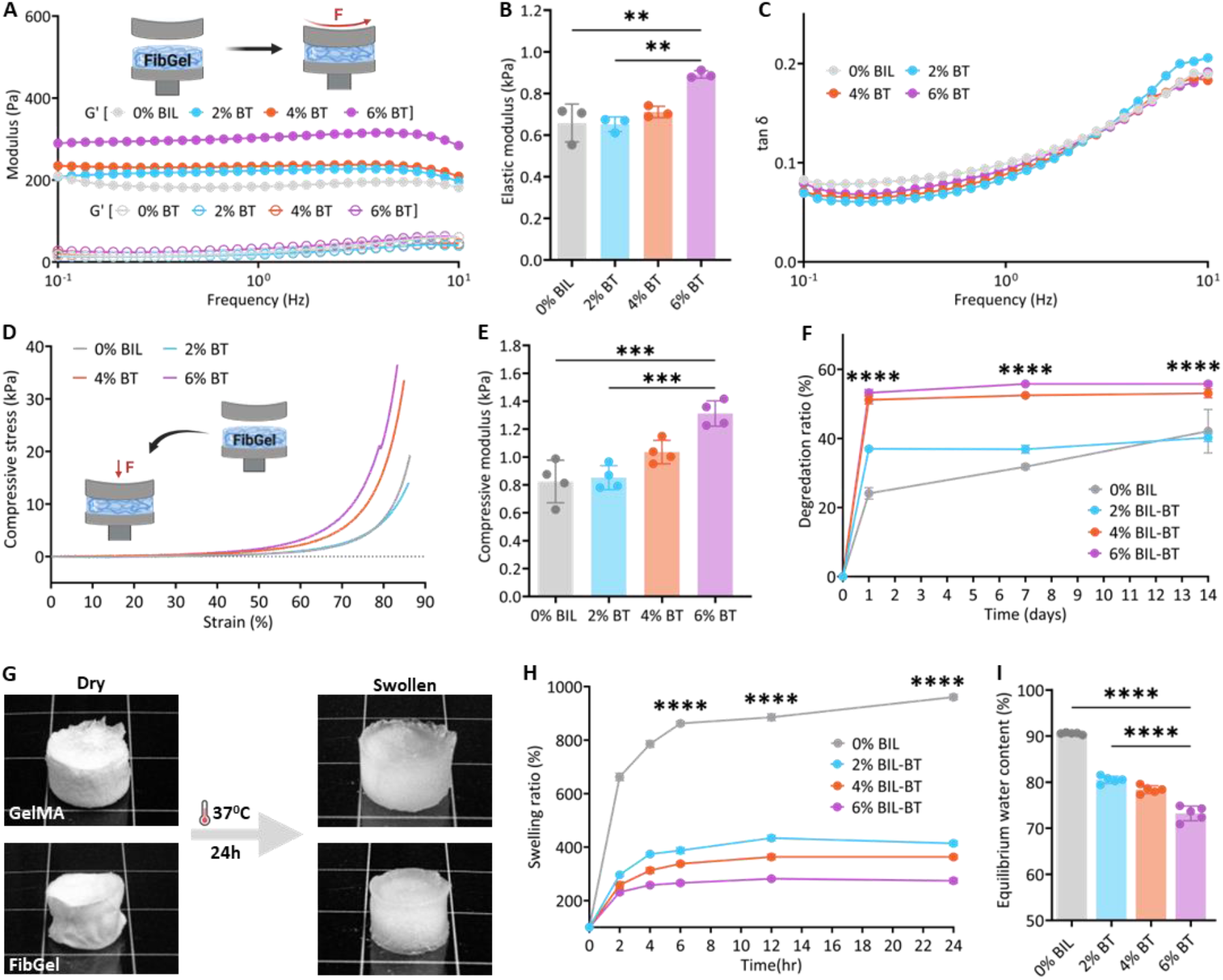
Mechanical, degradation, and swelling profiles of FiBGel hydrogels. (A) Frequency sweeps (0.1– 10 Hz) of shear (G′) and loss (G″) moduli for bioionic liquid (BIL)-conjugated fish gelatin methacryloyl (GelMA) hydrogels (FiBGel) prepared with 2, 4, or 6% (w/v) choline bitartrate (ChBT) and compared against 7% (w/v) GelMA controls (0% BIL). All hydrogel formulations were cured via 405 nm irradiation with LAP (lithium phenyl-2,4,6-trimethylbenzoylphosphinate, 0.25% w/v). (B) Elastic modulus deduced around 1 Hz increases with BIL concentration. (C) Frequency-dependent loss tangent (tan δ) profiles across the FiBGel hydrogels. (D) Unconfined compression stress–strain curves for FiBGel demonstrate pronounced, concentration-dependent stiffening. (E) Compressive modulus calculated from the initial strain region summarizes these stiffening trends. (F) Cumulative mass loss over 14 days in PBS (37 °C, pH 7.4) for FiBGel demonstrates degradation profiles dependent on BIL concentration. (G) Photographs of lyophilized discs (Dry) and the same samples after 24 h equilibration in PBS (37 °C, pH 7.4) (Swollen), illustrating the greater volumetric expansion of GelMA controls relative to FiBGel formulations. (H) Swelling ratio versus time for FiBGel hydrogels. Gels reach equilibrium swelling within 6 h, with higher BIL content yielding lower swelling ratios. (I) Equilibrium water content of FiBGel hydrogels formulated with 0, 2, 4, or 6% (w/v) ChBT. Values represent mean ± SEM, n = 3–5. Statistical significance via one-way ANOVA followed by Tukey’s post-hoc correction (ns: not significant, *p ≤ 0.05, **p ≤ 0.01, ****p ≤ 0.0001).

Uniaxial compression testing further probed the bulk response of the hydrated networks to normal loads, capturing the coupled solid-matrix deformation and fluid exudation (Figure 2D). All samples displayed the canonical “J-shaped” stress-strain profile characteristic of soft hydrogels[47,48], with a compliant toe (<10% strain) that transitioned to strain-stiffening beyond ∼ 20% strain. Compressive moduli deduced from the linear elastic region accentuate the unique structural influence of the bitartrate counter-ion. While GelMA controls registered 0.82 ± 0.15 kPa, ChBT incorporation resulted in only a modest increase to 1.31 ± 0.09 kPa at the maximum 6% (w/v) loading (Figure 2E). These mechanical profiles confirm that even at maximal ionic loading, FiBGel remains anchored within the optimal stiffness window (E ≤ 1.5 kPa) for neural tissue engineering.

A delicate balance between scaffold resorption and tissue formation is essential for brain tissue regeneration, where axonal extension and synaptic maturation require weeks to months. Gravimetric tracking in PBS (37 °C) confirmed that GelMA controls degraded gradually, registering a 24 ± 3% mass loss on Day 1 that progressed to 39 ± 6% by Day 14 (Figure 2F). In contrast, FiBGel displayed a distinct biphasic degradation profile. Day 1 mass loss increased with ionic liquid loading, reaching 38 ± 2% (2% w/v), 51 ± 2% (4% w/v), and 53 ± 2% (6% w/v). However, subsequent degradation plateaued, with mass loss advancing only marginally to ∼40–57% by Day 14. This profile indicates that after the rapid elution of uncrosslinked components, the remaining hydrogel network maintains significant structural robustness suitable for supporting tissue integration.

Gravimetric assays on lyophilized disks revealed a pronounced, concentration-dependent divergence in swelling ratio (Figure 2G–2I). The unmodified GelMA control absorbed ∼9.4 times its dry weight at 24 h, consistent with the 600–1000% range typically reported for GelMA hydrogels[49]. Introducing BIL side chains markedly suppressed this response and improved stability. FiBGel hydrogels stabilized at swelling ratios of 414 ± 21% (2% w/v), 363 ± 18% (4% w/v), and 274 ± 22% (6% w/v), reaching ≥ 95% of their plateau within 6 h(Figure 2H). This concentration-dependent stability dictates the equilibrium water content (EWC) of the final construct (Figure 2I). Whereas GelMA remained highly hydrated at 90.6 ± 0.3%, FiBGel formulations contracted to an EWC range of 72–81%. This interval precisely brackets the 78–82% water fraction characteristic of adult cortical gray matter [50,51], ensuring the implant matches the native tissue’s aqueous environment. Complementary dimensional tracking (Figure S5) confirmed that ChBT incorporation also curtails lateral expansion; while GelMA disks expanded to ∼122% of their initial area, FiBGel variants plateaued around 115% within 2 h, demonstrating superior retention of initial geometry for precise intracortical fitting.

### 2.4. Cytocompatibility and morphology of encapsulated neural stem cells

Cell viability and metabolic activity were evaluated to determine if BIL incorporation compromises cytocompatibility. To this end, iPSC-derived human neural stem cells (i-HNSCs) were encapsulated at 4 × 10⁶ cells mL^-1^ within the hydrogel network (Figure 3A). Embedding cells in this 3D environment reproduces the compliant, hydrated milieu of the brain parenchyma, avoiding the artificial adhesion cues typical of 2D cultures[52]. Live/Dead imaging from 7 days post-encapsulation (dpe) revealed high cell viability across all FiBGel formulations, with little to no ethidium-homodimer uptake detected even at the maximum 6% (w/v) loading (Figure 3B & S6). Metabolic tracking via PrestoBlue assay corroborated this viability profile (Figure 3C–E). While the metabolic baseline at 1 dpe appeared modestly reduced in ChBT groups (∼0.22–0.35 a.u.) compared to GelMA controls (∼0.38 a.u.), activity recovered over time. By 14 dpe, the 2% and 4% (w/v) ChBT were statistically indistinguishable from the GelMA control. This confirms that the incorporated bioionic liquids are intrinsically non-toxic and support sustained metabolic proliferation.

**Figure 3.**
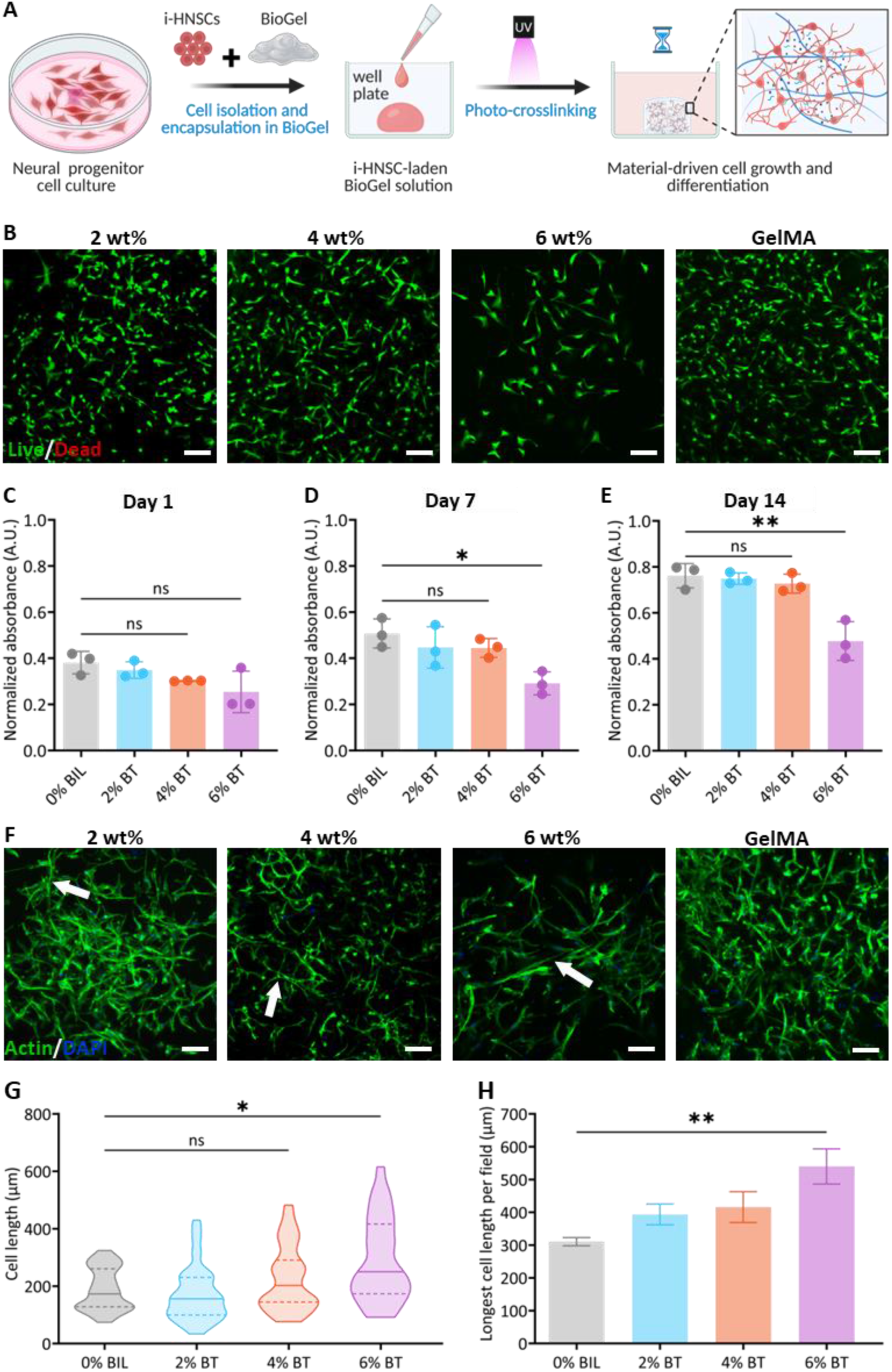
Viability and morphological evalualtion of i-HNSCs in FiBGel matrices. (A) Schema of the experimental workflow for encapsulating iPSC-derived human neural stem cells (i-HNSCs) in bioionic liquid (BIL)-conjugated fish gelatin methacryloyl (GelMA) hydrogels (FiBGel), followed by photocrosslinking and culture. (B) Representative confocal micrographs of i-HNSC viability at 14 days post-encapsulation (dpe), labeled with calcein-AM (viable cells, green) and ethidium homodimer-1 (dead cells, red). Cells were cultured in hydrogels composed of 7% (w/v) GelMA functionalized with 0, 2, 4, or 6% (w/v) choline bitartrate (ChBT). The 0% BIL formulation serves as the control, and all hydrogel formulations were photocrosslinked using LAP (lithium phenyl-2,4,6-trimethylbenzoylphosphinate, 0.25% w/v). Scale bars: 100 µm. (C–E) Quantification of metabolic activity via PrestoBlue assay at 1 dpe (C), 7 dpe (D), and 14 dpe (E), demonstrating the dependence of cell proliferation on BIL concentration over time. (F) Confocal images of F-actin (phalloidin, green) and nuclei (DAPI, blue) at 14 dpe within FiBGel formulations. i-HNSCs progressively adopt elongated morphologies as ChBT content increases (white arrows). Scale bars: 100 µm. (G) Distribution of cell lengths across all hydrogel formulations. (H) Longest cell length per field, highlighting the impact of ChBT concentration on maximal elongation. Values represent mean ± SEM. Statistical significance via one-way ANOVA followed by Tukey’s post-hoc correction. (ns: not significant, *p ≤ 0.05, **p ≤ 0.01).

Neuromimetic matrices drive neuronal lineage commitment by enhancing cytoskeletal tension generation and lowering the energetic barrier for the dynamic F-actin remodeling that underpins growth cone advancement[53]. To probe how the electroactive microenvironment modulates these morphodynamic cues, i-HNSCs were visualized via phalloidin/DAPI staining. At 7 dpe, slender, interconnected actin filaments were observed in FiBGel (Figure S7). By 14 dpe, this reorganization evolved into extensive, bipolar F-actin networks. Specifically, hydrogels with 6% (w/v) ChBT exhibited elongated actin cables and neurite-like projections spanning hundreds of micrometers (Figure 3F, white arrows). Quantitative morphological analysis revealed a concentration-dependent increase in median process length (Figure 3G). Notably, the 6% (w/v) formulation induced a ∼1.7-fold increase in the maximum neurite extension relative to unfunctionalized GelMA controls (Figure 3H). This extensive elongation indicates that the electroactive FiBGel niche actively drives cytoskeletal tension generation and morphological maturation.

### 2.5. Neurogenic Differentiation and Phenotypic Maturation

Physical and electrochemical cues inherent to the extracellular matrix can independently direct lineage commitment, potentially bypassing the need for exogenous inductive factors. We interrogated this by culturing encapsulated i-HNSCs in proliferation medium devoid of differentiation factors for 28 days within hydrogels containing 0, 2, 4, or 6% (w/v) choline bitartrate (designated GelMA, FiBGel-2, FiBGel-4, and FiBGel-6, respectively). Immunostaining unveiled a robust, dose-dependent upregulation of the early neuronal marker β3-tubulin (β3T) (Figure 4A). The FiBGel-6 formulation triggered an 8-fold increase in β3T+ cells (63.5%) relative to GelMA controls (8.2%), confirming that material properties alone effectively prime the progenitor pool. When combined with standard biochemical induction at 7 dpe, these matrix cues synergized to expedite phenotypic maturation. Confocal micrographs (Figure 4B, top) reveal progressively higher MAP2+ cells with increasing BIL content, while the corresponding quantification (Figure 4B, bottom) shows a progressive shift from “β3T-only” toward “MAP2 + β3T” co-expression. The population of dual-positive neurons (MAP2+/β3T+) expanded from 56.1% in GelMA to 90.3% in FiBGel-6. Consequently, the electroactive niche converted nearly the entire progenitor pool into functionally mature neurons by day 21 post-differentiation, significantly accelerating the maturation timeline compared to unfunctionalized hydrogels.

**Figure 4.**
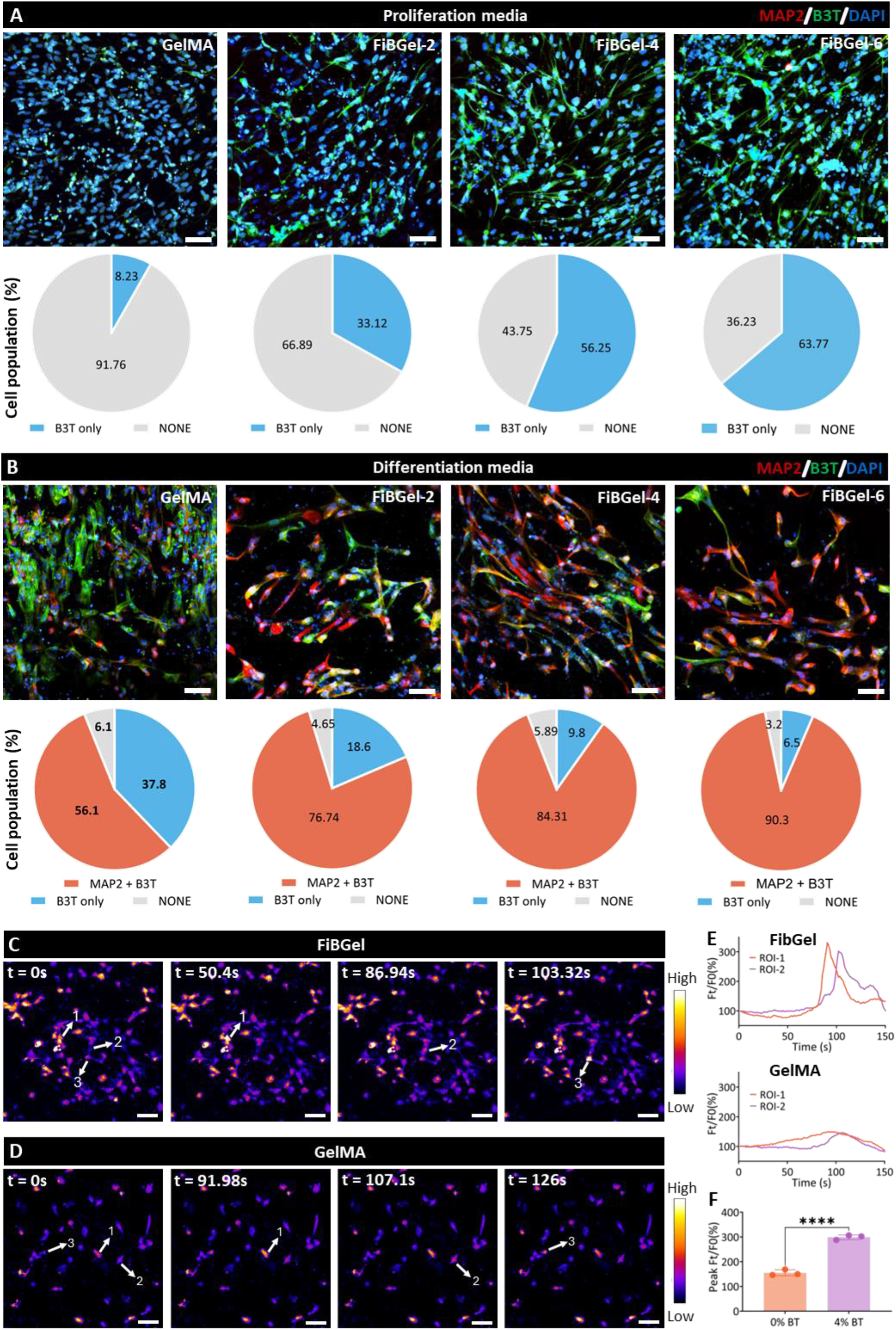
Neuronal differentiation and calcium activity of i-HNSCs in FiBGel hydrogels. (A) Confocal micrographs and accompanying population pie charts for iPSC-derived human neural stem cells (i-HNSCs) at 28 days post-encapsulation (dpe) in proliferation medium within bioionic liquid (BIL)-conjugated fish gelatin methacryloyl (GelMA) hydrogels (FiBGel), containing 2, 4, or 6% (w/v) choline bitartrate (ChBT) (labelled FiBGel-2, FiBGel-4, and FiBGel-6, respectively). Cells were stained for the early neuronal marker β3-tubulin (B3T, green) and nuclei (DAPI, blue); the fraction of B3T+ cells rises with increasing ChBT content. (B) Parallel cultures in differentiation medium (added at 7 dpe) were stained for both B3T (green) and microtubule associated protein-2 (MAP2, red). Co-expression of these markers increased significantly with ChBT concentration, demonstrating accelerated neuronal maturation. (C, D) Time-lapse calcium imaging of networks formed in FiBGel-4 (C) versus a GelMA control (D). Pseudocolour heat maps depict spontaneous intracellular Ca²⁺ transients; arrows highlight representative active cells. (E) Normalized fluorescence traces (F/F₀) from different regions of interest reveal higher amplitude and more frequent calcium spikes in FiBGel-4 relative to the GelMA control. (F) Quantification of peak fluorescence intensity confirms significantly enhanced functional activity in networks formed within FiBGel-4 hydrogels. All hydrogels were composed of 7% (w/v) GelMA and photocrosslinked with 0.25% (w/v) lithium phenyl-2,4,6-trimethylbenzoylphosphinate (LAP). Scale bars: 50 µm. Values represent mean ± SEM. Statistical significance via a two-tailed unpaired t-test (**** p ≤ 0.0001).

Functional network connectivity was subsequently interrogated using Fluo-8® AM calcium imaging at 21 days post-differentiation. This assessment was performed to capture intracellular calcium dynamics without external stimulation, thereby highlighting intrinsic network activity. We selected FiBGel-4 for these tests to balance neuromimetic properties, cytocompatibility and neurogenic potential. Pseudocolour time-lapse frames highlight prominent, high-amplitude Ca²⁺ transients propagating across multiple cells in the FiBGel matrix (Figure 4C, arrows 1–3), a striking contrast to the sparse, low-intensity activity observed in GelMA controls (Figure 4D). Normalized fluorescence traces quantified this enhancement, revealing that peak intensities (F_t_/F_0_) rose from 152 ± 18% in controls to 296 ± 21% in FiBGel (Figure 4E). This ∼1.9-fold amplification in spontaneous excitability confirms that the electroactive microenvironment effectively lowers the excitation threshold, driving the robust synaptic signaling and spontaneous neuronal firing essential for functional tissue integration.

### 2.6. 3D Bioprinting of cell-Laden constructs

While extrusion-based bioprinting remains prevalent, its application to soft hydrogels is often constrained by shear stresses that compromise cell viability[54]. Conversely, nozzle-free vat polymerization via digital light processing (DLP) eliminates these shear forces and decouples print duration from construct complexity, characteristics that are particularly advantageous for fabricating soft, brain-like tissues with high cell survival rates[55]. Leveraging this, we utilized DLP to fabricate complex neural architectures using the FiBGel-4 formulation. Initial acellular parameter sweeps modulating photoabsorber concentration and exposure time culminated in the high fidelity production of anatomical brain models at ∼18.8 mW/cm² irradiance and 50 μm layer height (Figure S8 & S9A) (More details in Supplementary Note 2). Transitioning to cell encapsulated bioinks, we removed cytotoxic photoabsorbers and utilized the culture medium’s intrinsic attenuation to cure 100 μm layers at a reduced energy density (∼10.96 mW/cm²), thereby preserving the delicate stem cell payload. Biological assessment of these i-HNSC encapsulated constructs at Day 7 revealed extensive axonal projections and high viability (Figure S9B), a trajectory that accelerated by Day 14 to exhibit robust, interconnected neural networks (Figure S9C). This successful adaptation confirms the capacity of the optimized bioink to support scalable, spatially defined tissue engineering without compromising the cytocompatibility advantages of the soft, electroactive matrix.

### 2.7. Pro-angiogenic activity of bioionic hydrogels

The metabolically demanding environment of developing neural tissue necessitates rapid vascularization to deliver oxygen and neurotrophic factors. We evaluated the capacity of the electroactive FiBGel matrix to support 3D tubulogenesis by encapsulating human umbilical vein endothelial cells (HUVECs) within hydrogels containing varying choline bitartrate concentrations (0–6% w/v). Phase-contrast imaging at 14 days post-encapsulation revealed the progressive self-assembly of capillary-like structures, which became increasingly dense and interconnected with higher choline bitartrate loading (Figure S10). Immunofluorescence analysis of endothelial markers CD31 and VE-cadherin confirmed this robust network formation (Figure 5A). Quantitative morphometry demonstrated a significant, dose-dependent escalation in both branch density (Figure 5B) and junctional frequency (Figure 5C), with a distinct inflection point observed between the FiBGel-2 and FiBGel-4 formulations. This response suggests that a specific threshold of bioionic concentration is required to markedly accelerate endothelial organization.

**Figure 5.**
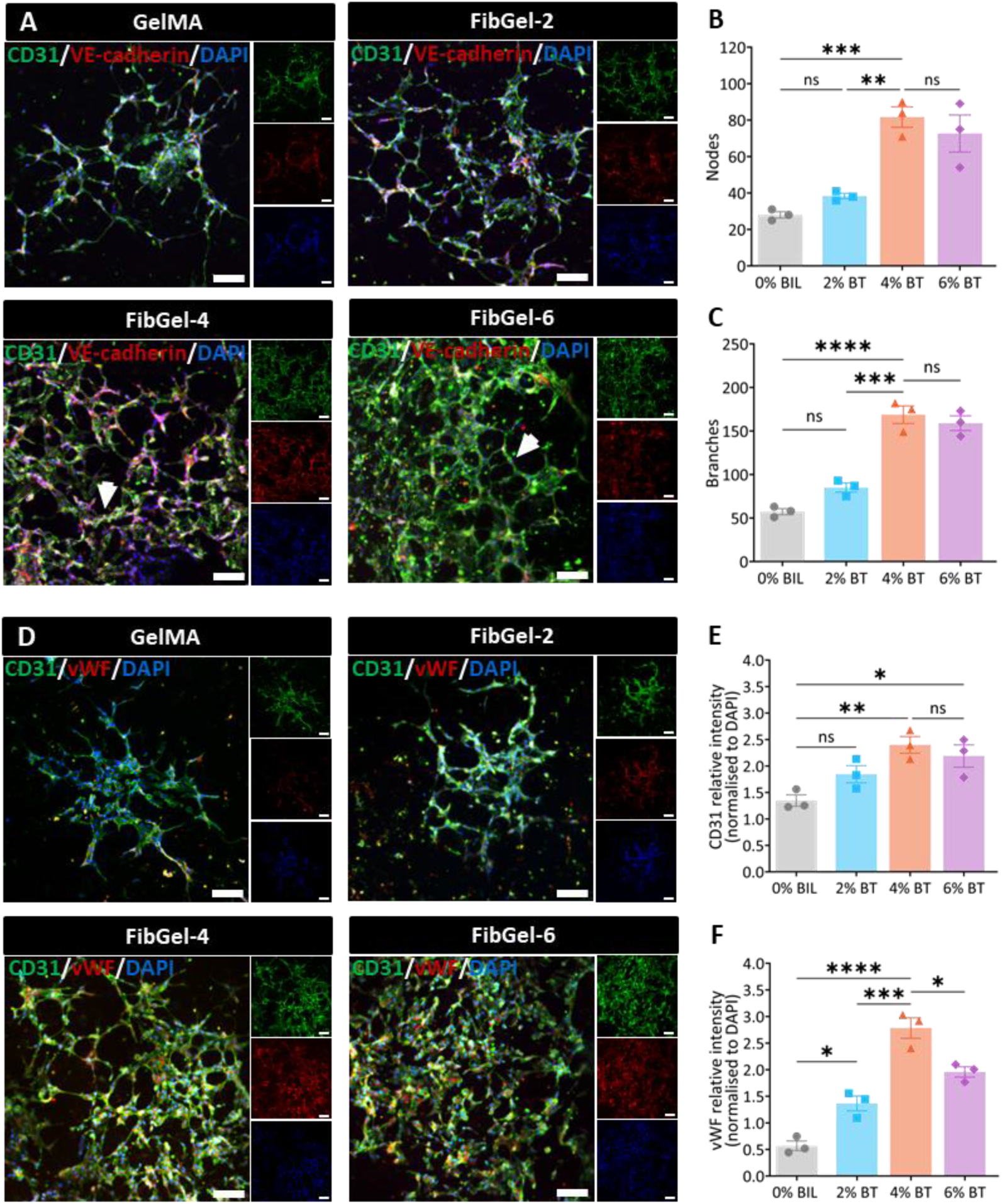
Angiogenic response of HUVECs in FiBGel hydrogels. (A) Confocal images of human umbilical vein endothelial cells (HUVECs) cultured for 21 d in 7% (w/v) fish gelatin methacryloyl (GelMA) or in bioionic liquid-conjugated GelMA (FiBGel) containing 2, 4, or 6% (w/v) choline bitartrate (ChBT) (labelled FiBGel-2, FiBGel-4, and FiBGel-6, respectively). Cells were stained for CD31 (green), vascular endothelial cadherin (VE-cadherin, red), and nuclei (DAPI, blue). Increasing ChBT content promoted denser, highly branched microvascular networks (white arrowheads). (B, C) Quantification of network complexity via number of branch nodes (B) and branches per field (C). Both metrics rose significantly at 4 and 6% (w/v) ChBT relative to GelMA controls, indicating enhanced tubulogenesis. (D) Representative images of CD31 (green) and von Willebrand factor (vWF, red) staining illustrate endothelial maturation. (E, F) Mean fluorescence intensity, normalized to DAPI, for CD31 (E) and vWF (F) increases with ChBT concentration, confirming elevated endothelial marker expression. Scale bars: 100 µm. Values represent mean ± SEM(n=4). Statistical significance via one-way ANOVA with Tukey post-hoc correction (ns: not significant, *p ≤ 0.05, **p ≤ 0.01, ***p ≤ 0.001, ****p ≤ 0.0001).

Endothelial maturation was further corroborated by the expression of von Willebrand factor (vWF), a glycoprotein essential for hemostasis and vascular integrity. Confocal imaging revealed that HUVECs within FiBGel constructs matured into a stable endothelial phenotype characterized by elevated vWF and CD31 co-expression (Figure 5D). Normalized fluorescence intensity analysis indicated that marker expression peaked within the FiBGel-4 formulation (Figure 5E & 5F), suggesting an optimal bio-ionic window for vascular development. While FiBGel −6 supported extensive branching, the slight plateau in maturation markers compared to the 4% formulation implies a limit where osmotic or signaling effects may become saturating. Indeed, FiBGel-4 emerges as the superior candidate, balancing extensive network complexity with robust phenotypic maturation.

### 2.8. Biocompatibility and neuroinflammatory modulation

Evaluating the neuroimmune response constitutes a fundamental requirement for validating biomaterials intended for central nervous system applications, where glial cells function as primary sentinels coordinating inflammatory cascades that dictate therapeutic outcomes[56,57]. Stereotactic intracortical injection into C57BL/6J mice was employed to interrogate the local and regional neuroimmune response (Figure 6A), comparing FiBGel against vehicle groups with non-injected mice serving as sham controls. Immunohistochemical profiling at post-injection day (PID) 7 revealed a distinct modulation of microglial surveillance in animals treated with FiBGel (Figure 6B). Staining for the ionized calcium-binding adaptor molecule 1 (Iba1) demonstrated a marked reduction of microglial activation compared to vehicle controls. Quantitative regional analysis confirmed this immunosuppressive potency, registering significant reductions in Iba1 expression within both the ipsilateral cortex (p < 0.01) and the underlying hippocampus (p < 0.001) (Figure 6C). This indicates that the hydrogel actively dampens the acute inflammatory cascade rather than merely occupying space as an inert filler.

**Figure 6.**
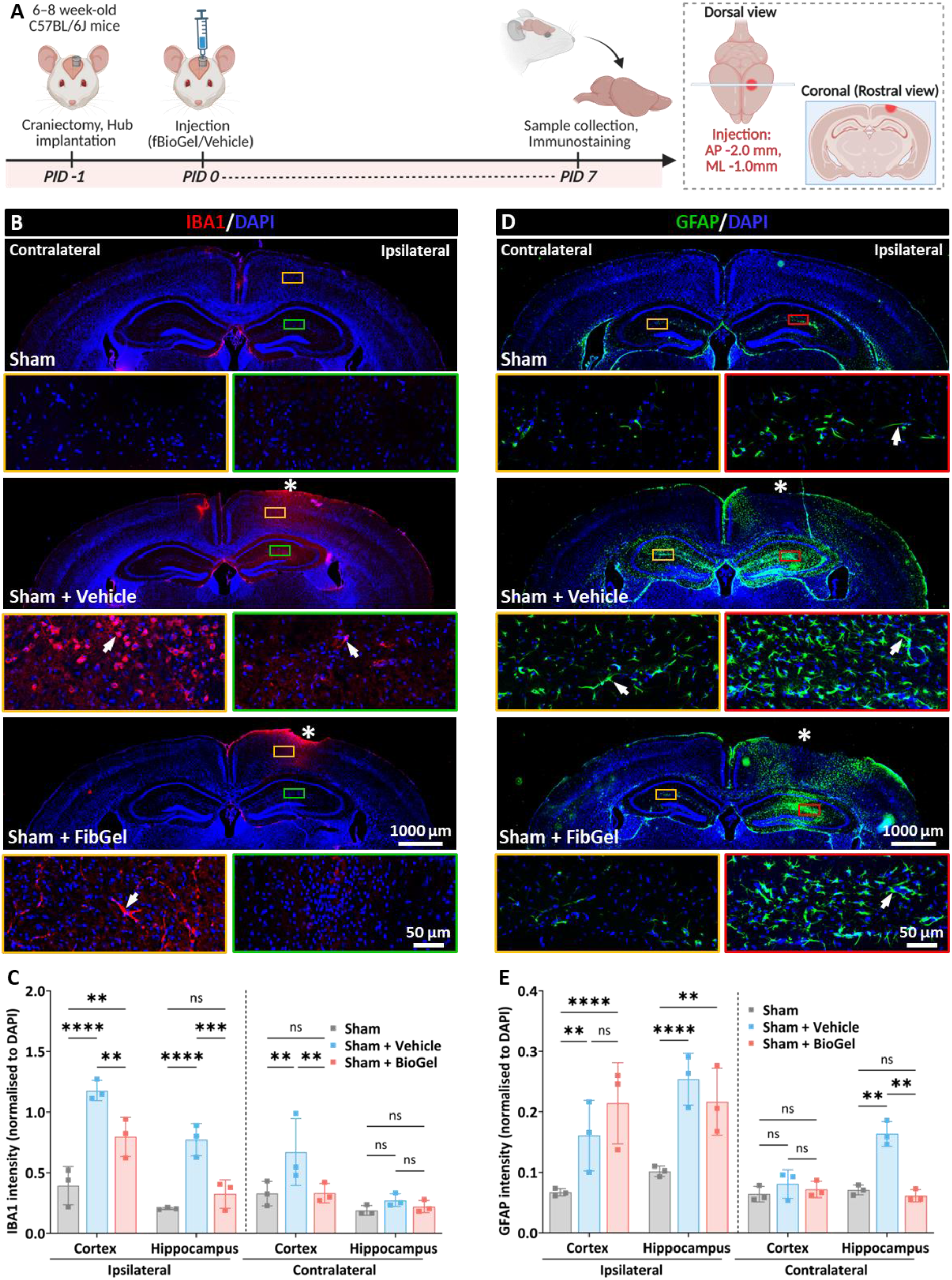
Biocompatibility of FiBGel following stereotactic injection in the cortex. (A) Experimental timeline. Mice underwent a left hemisphere craniectomy, and saline (Vehicle) or bioionic liquid– conjugated fish gelatin methacryloyl (GelMA) hydrogel (FiBGel) was injected stereotactically (AP –2.0 mm, ML –1.0 mm relative to bregma). Animals were sacrificed and brains collected at post-injection day (PID) 7. (B, D) Coronal sections showing the injection site (asterisk) stained for microglia (Iba1, red) (B) or reactive astrocytes (GFAP, green) (D) with DAPI-labelled nuclei (blue). Sham animals serve as surgical controls, and contralateral hemispheres provide internal references. White arrows highlight representative activated glial cells. (C, E) Quantification of Iba1 (C) and GFAP (E) fluorescence, normalized to DAPI, in the ipsilateral and contralateral regions. Scale bars: low magnification, 1000 µm; high magnification, 50 µm. Values represent mean ± SEM. Statistical significance via two-way ANOVA with Tukey’s post-hoc correction (ns, not significant; **p ≤ 0.01, ***p ≤ 0.001, ****p ≤ 0.0001). AP, anteroposterior; DV, dorsoventral; Iba1, ionised calcium-binding adaptor molecule 1; GFAP, glial fibrillary acidic protein; ML, mediolateral.

Astrocytic reactivity, assessed via GFAP expression, exhibited a more complex, spatially distributed response pattern (Figure 6D). While the cortex displayed non-significant reductions in normalized GFAP fluorescence, analysis of the hippocampal circuitry unveiled a salient distal effect. Notably, FiBGel administration elicited a significant suppression of astrocytic activation within the contralateral hippocampus (p < 0.01) relative to vehicle groups (Figure 6E). Furthermore, analysis of the lesion area corroborated the material’s safety; the cortical damage remained statistically indistinguishable from vehicle controls (Figure S12A). Collectively, these findings demonstrate that FiBGel exerts a potent, spatially extensive immunomodulatory influence, attenuating glial reactivity across neural networks beyond the implantation site.

### 2.9. Therapeutic efficacy in traumatic brain injury

TBI is a major global contributor to death and disability, affecting around 69 million individuals annually[1]. TBI triggers complex pathophysiological cascades that evolve from the primary mechanical insult into secondary injury mechanisms characterized by blood-brain barrier disruption, excitotoxicity, and sustained glial scarring. To model these pathological signatures with high translational fidelity, we utilized a lateral fluid percussion injury (FPI) model (Figure S11) at a moderate severity (1.5–1.7 atm), which produces reproducible neurological deficits and tissue damage patterns mirroring clinical outcomes[58–61]. Immediately following injury, animals received intracortical injections of vehicle, GelMA, or FiBGel, and coronal sections spanning the impact site were analyzed at 7 days post injury (dpi) to capture the peak window of astrocytic reactivity (Figure 7A) [62]. Non-injected FPI groups (Figure S13) were excluded to isolate material effects, as needle insertion and injection exacerbate the local inflammatory baseline[63,64]. While microglial activation (Iba1) remained comparable across groups, indicating that the subacute microglial burden is largely driven by the primary insult (Figure 7B & 7C), astrocytic responses were profoundly modulated. Immunofluorescence imaging revealed markedly reduced GFAP expression in the ipsilateral cortex of FiBGel-treated animals compared to controls (Figure 7D). Quantitative analysis confirmed at least a 2-fold reduction in astrogliosis (p < 0.0001) compared to GelMA and vehicle groups, an effect that notably extended to the contralateral hippocampus (Figure 7E). This echoes our biocompatibility assessment and indicates systemic astrogliotic modulation that extends beyond the primary injury epicenter.

**Figure 7.**
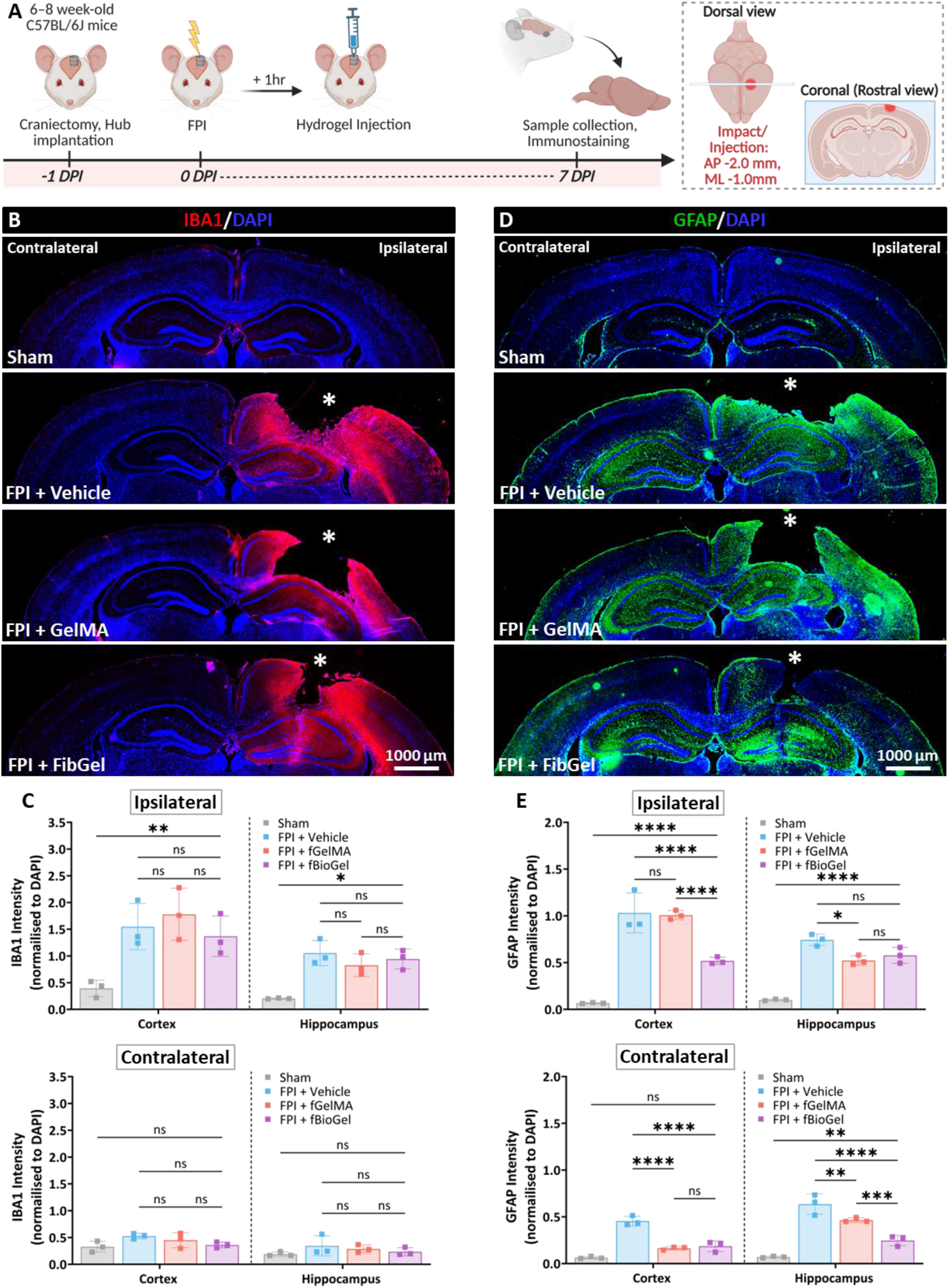
Neuroprotective effects of FiBGel in a fluid percussion injury model. (A) Schematic of experimental timeline. Mice underwent craniectomy over the left parietal cortex and hub implantation 24 h before moderate fluid percussion injury (FPI). Following FPI, saline (Vehicle), fish gelatin methacryloyl (GelMA), or bioionic liquid–conjugated GelMA (FiBGel) was injected stereotactically into the ipsilateral cortex (AP –2.0 mm, ML –1.0 mm from bregma). Brains were harvested at 7 days post-injury (dpi) for histological analysis. (B, D) Representative coronal sections through the impact site (asterisk) immunostained for activated microglia (Iba1, red) (B) or reactive astrocytes (GFAP, green) (D) with nuclear counterstaining (DAPI, blue). Experimental groups include Sham (surgical control), FPI + Vehicle (injection control), FPI + GelMA, and FPI + FiBGel, comparing ipsilateral and contralateral hemispheres. White arrows indicate representative activated glial cells. (C, E) Quantification of Iba1 (C) and GFAP (E) immunofluorescence intensity normalized to DAPI, in ipsilateral and contralateral regions. Scale bars: 1000 µm. Values represent mean ± SEM. Statistical significance via one-way ANOVA with Tukey’s post-hoc correction (ns: not significant, *p ≤ 0.05, **p ≤ 0.01, ***p ≤ 0.001, ****p ≤ 0.0001). AP, anteroposterior; DV, dorsoventral; Iba1, ionised calcium-binding adaptor molecule 1; GFAP, glial fibrillary acidic protein; ML, mediolateral.

Beyond glial modulation, FiBGel treatment orchestrated a robust, region-specific vascular remodeling response. Immunohistochemical staining for the endothelial marker (CD31) at 7 dpi revealed dense perilesional microvessel networks in FiBGel-treated mice, contrasting with the sparse vasculature observed in vehicle and GelMA groups (Figure 8A). Quantification confirmed a significant increase in CD31 intensity within the ipsilateral cortex relative to vehicle (p < 0.01) and GelMA (p < 0.05), with a smaller yet significant elevation in the contralateral cortex compared to GelMA (p < 0.05) (Figure 8B & 8C). While hippocampal values trended upward without reaching significance, the preferential cortical revascularization suggests that FiBGel establishes a pro-regenerative niche. By concurrently limiting glial scarring and rebuilding the microvasculature, the material effectively counteracts the capillary rarefaction and hypoperfusion that exacerbate metabolic stress in acute TBI[65]. This dual therapeutic effect culminated in superior tissue preservation as FiBGel treatment significantly reduced the cortical lesion area relative to both GelMA and vehicle groups (p < 0.0001, Figure S12B), confirming that the hydrogel actively limits secondary tissue damage.

**Figure 8.**
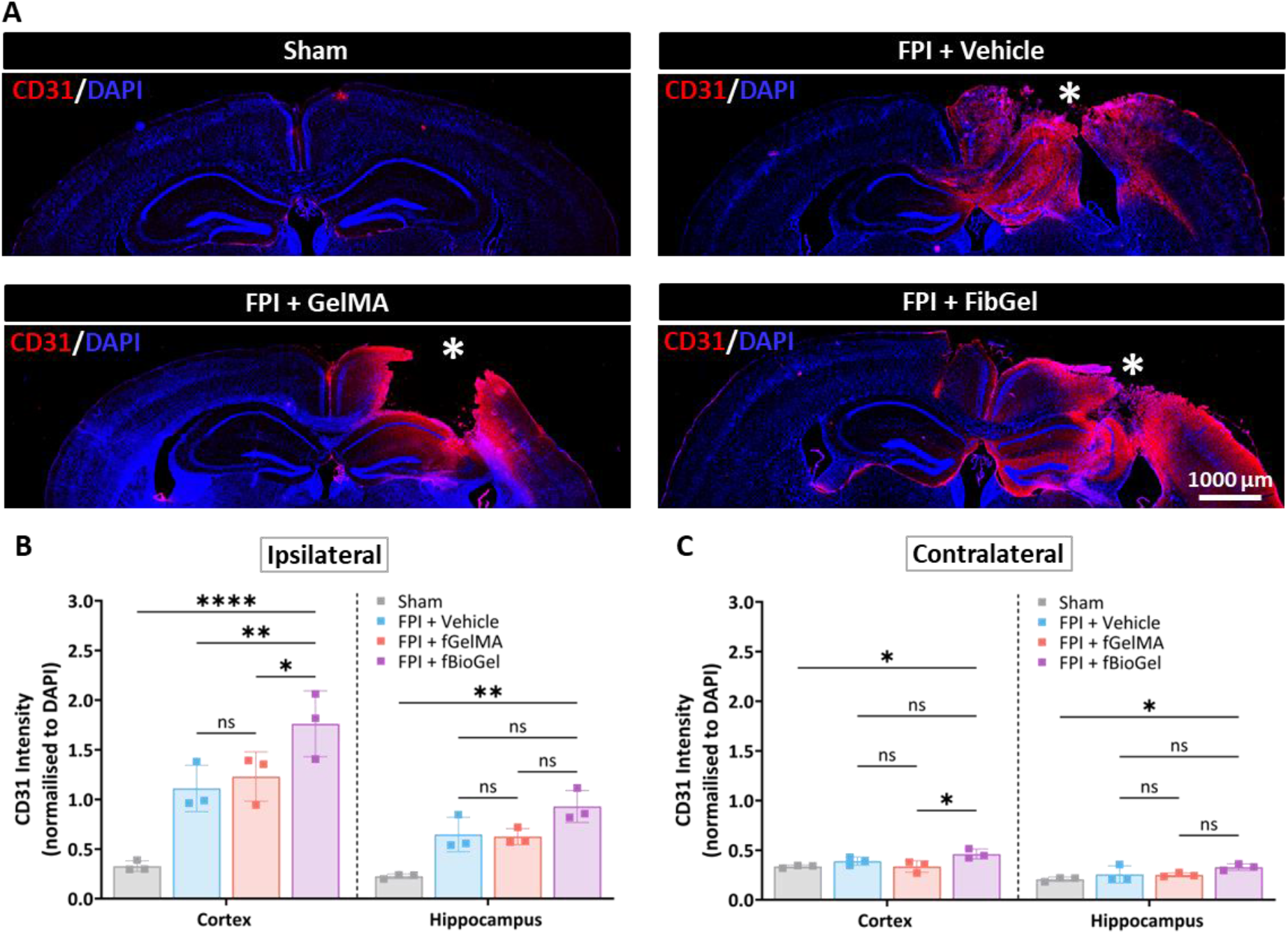
Angiogenic potential of FiBGel in a fluid percussion injury model. (A) Representative coronal brain sections at 7 days post-injury (dpi) immunostained for endothelial cells (CD31, red) with nuclear counterstaining (DAPI, blue). Experimental groups include Sham (surgical control), FPI + Vehicle (injection control), FPI + fish gelatin methacryloyl (GelMA), and FPI + bioionic liquid–conjugated GelMA (FiBGel). The impact site is marked with an asterisk (*). Images compare ipsilateral and contralateral hemispheres, demonstrating enhanced vascularization in FiBGel-treated animals relative to other groups. (B, C) Quantification of CD31 immunofluorescence intensity normalized to DAPI, in the ipsilateral (B) and contralateral (C) cortex and hippocampus. FiBGel treatment shows enhanced angiogenic response compared to Vehicle and GelMA controls. Scale bars: 1000 μm. Data represents mean ± SEM; statistical analysis via one-way ANOVA with Tukey’s post-hoc comparisons (ns: not significant, *p ≤ 0.05, **p ≤ 0.01, ****p ≤ 0.0001).

## 3. Discussion

The central paradox of neural bioelectronics is that the most robust electrical performance is often delivered by stiff materials that compromise cytocompatibility and exacerbate neuroimmune activation[57,66]. Conventional strategies often rely on percolation networks of rigid carbon or metallic fillers, which elevate composite stiffness well above the native brain regime and introduce formulation-dependent cytotoxicity risks [14,15]. We circumvent this paradigm by covalently tethering choline-based bioionic liquids onto a fish gelatin backbone, creating a “FiBGel” platform that harmonizes physiological conductivity (0.25–0.68 S m⁻¹) with the ultrasoft mechanics (0.6–0.9 kPa) and brain-matched water content (72–81%) essential for neural integration. Notably, the reported conductivities span reported cortical grey matter baselines (∼0.2–0.6 S m⁻¹)[67,68], ensuring seamless bioelectric coupling. Unlike electron-conducting scaffolds, our approach leverages mobile ions within a hydrated polymer network, aligning with the electrochemical basis of native neural signaling. The preservation of neurocompliant mechanics, even at high ionic loading, is governed by the specific chemistry of the bitartrate counter-ion. Functioning as a bulky steric spacer, the bitartrate moiety prevents the dense polymer chain aggregation that typically drives stiffening in functionalized networks [46]. This molecular configuration allows the hydrogel to sustain a viscoelastic profile matching native brain tissue [74–76], effectively mitigating the stiffness-conductivity trade-off that constrains conventional neural interfaces.

The biophysical and electrochemical fidelity establishes a highly cytocompatible and instructive niche capable of biasing neuronal specification independent of exogenous factors. Notably, we detected negligible cytotoxicity even at 6% (w/v) ionic liquid loading, standing in sharp contrast to carbon-based fillers that often elicit oxidative stress at concentrations as low as 0.002% (w/v)[11,12]. Upon biochemical induction, FiBGel drove nearly complete neuronal conversion, yielding ∼90% MAP2+ neurons with amplified calcium dynamics, an outcome that suggests a convergence of three material-derived cues. First, the maintenance of ultrasoft mechanics (<1 kPa) is consistent with reduced YAP/TAZ mechanotransduction, a pathway known to bias stem cell fate toward neuronal lineages on compliant substrates [69,70]. Second, the enhanced bulk conductivity likely creates a permissive ionic milieu that supports efficient redistribution of endogenous ionic currents, facilitating activity-dependent network assembly [71]. Third, the grafted cholinium moieties may engage cholinergic signaling such as α7-nicotinic acetylcholine receptor (α7-nAChR), initiating cascades that reinforce neuronal identity [72]. These mechano-electrochemical cues first prime the progenitor pool, evidenced by an eight-fold increase in early neuronal commitment (β3T) without exogenous factors, and subsequently accelerate phenotypic maturation. Indeed, by potentiating endogenous bioelectric signaling and reinforcing neurogenic cues via material features, the bioengineered niche effectively compresses the maturation timeline of human neuronal cells.

Effective neural tissue repair demands biomaterials that transcend the role of passive cell carriers to actively intervene in the secondary injury cascades that drive chronic neurodegeneration. Our results suggest that FiBGel functions as a bioresponsive immunomodulator capable of reshaping the injury microenvironment, as evidenced by the profound attenuation of neuroglial activation and significant reduction in cortical lesion expansion. This therapeutic effect is likely mediated by the bioactive cholinium component, which acts as a precursor for acetylcholine to activate the cholinergic anti-inflammatory pathway via α7-nAChR on glial cells [23,24]. By curtailing pro-inflammatory cytokine release, this signaling axis intercepts the inflammatory feedback loops propagating through transcallosal projections [73,74] underpinning the distinct bilateral suppression of glial reactivity. Complementing this biochemical signaling, the shape fidelity and modulus matching of the hydrogel likely minimize the mechanical stress on the host parenchyma, thereby reducing the release of damage-associated molecular patterns (DAMPs) that typically trigger sustained gliosis [75,76]. These biochemical and physical cues appear to temper glial activation and preserve tissue architecture, consequently mitigating necrotic cavity expansion typical of moderate TBI.

The restoration of a functional neurovascular unit stands as a prerequisite for neural tissue survival, as the temporal and spatial regulation of angiogenic stimuli determines whether regenerating vasculature supports functional repair or devolves into disorganized networks[77]. Extending our prior report of angiogenesis using BIL hydrogels in 2D cultures[33], our current findings indicate that FiBGel orchestrates a robust 3D vascular response driven by convergent material cues. Mechanistically, the ∼1 kPa modulus lies within the permissive stiffness window (1–6 kPa) that favors endothelial sprouting and the upregulation of tip cell genes including VEGFR2 and DLL4[78,79], while the ionic conductivity of the matrix likely potentiates electrochemical coupling via gap junctions[80]. This is further reinforced by choline release, which functions as a pharmacological agonist for α7 nicotinic receptors to upregulate endogenous VEGF expression[26,81]. In the context of TBI, this enhanced revascularization directly counters the acute capillary rarefaction and hypoperfusion that drive the post-traumatic metabolic crisis[65]. By restoring microvascular density, FiBGel likely improves oxygen–glucose delivery and accelerates clearance of excitotoxic by-products, thereby mitigating secondary neuronal loss. Consistent with recent evidence that microporous conductive hydrogels reduce scarring[82], these findings suggest that FiBGel establishes a self-reinforcing loop of vascular supply and neural survival extending from the injury core to the penumbra.

In conclusion, this work establishes a new paradigm for neural biomaterial design by demonstrating that bioionic liquid functionalization can simultaneously optimize electromechanical fidelity and biological performance within a single injectable platform. The convergence of tissue-matched mechanics, neuroinductive, anti-inflammatory, and pro-angiogenic properties renders FiBGel a versatile therapeutic for the complex pathophysiology of brain injury and neurodegenerative diseases. Moreover, the successful adaptation of the platform for DLP bioprinting opens new avenues for fabricating personalized neural tissue constructs and high-throughput disease models. Given the FDA-recognized safety of choline bitartrate as a dietary supplement and the demonstrated safety profile of the platform, FiBGel represents a highly translatable solution that could significantly advance minimally invasive neuroregenerative therapies and improve functional outcomes for patients with neurological disorders.

## 4. Methods

### 4.1. GelMA Synthesis

Fish gelatin methacryloyl (GelMA) was synthesized in a similar manner to porcine GelMA synthesis in our previous studies[28,29]. In brief, 10 g of cold-water fish skin gelatin (Sigma-Aldrich) was dissolved at 50 °C under an inert atmosphere with continuous stirring in 100 mL PBS (pH 7.5, ThermoFisher) until complete dissolution. 8 mL of methacrylic anhydride (Sigma-Aldrich) was added dropwise over 15 min, and the methacrylation proceeded for 2 h at 50 °C. The reaction mixture was diluted with warm PBS (100 mL, 50 °C) and stirred until homogeneous. The resulting GelMA solution was purified by dialysis (12–14 kDa MWCO tubing, Repligen) against deionized water at 50 °C for 5 days to eliminate unreacted methacrylic anhydride and reaction byproducts. The purified solution was frozen at −80°C and subsequently lyophilized for 72 h to yield a porous foam, which was stored at −20 °C until use.

### 4.2. Acrylated choline bitartrate Synthesis

Acrylated choline bitartrate (ChBT) was synthesized following a previously reported protocol [29] with slight modification. In brief, we dissolved 16 g of choline bitartrate (Sigma-Aldrich) in 20 mL of deionized water at 50 °C under constant stirring in a 50 mL round-bottom flask. Under a nitrogen atmosphere, 5 g of acrylic acid (1.1 mol eq, Sigma-Aldrich) was introduced dropwise. The reaction proceeded under 50 mbar reduced pressure for 24 h at 50 °C to continuously remove water byproducts and drive the equilibrium toward esterification. The reaction mixture was concentrated and purified by washing with 30 mL ethyl acetate (× 5) to remove unreacted acrylic acid and any non-covalently bound byproducts, discarding the organic phase each time. Residual solvent was removed by rotary evaporation, and the resulting product was frozen at −80°C and lyophilized for 24 h to yield the acrylated choline bitartrate, which was stored under inert atmosphere until use.

### 4.3. Fabrication of FiBGel Hydrogels

Bioionic liquid (BIL)-conjugated fish gelatin methacryloyl (FiBGel) hydrogels were fabricated using GelMA at 7% (w/v), lithium phenyl-2,4,6-trimethylbenzoylphosphinate (LAP, 0.25% w/v, Sigma-Aldrich) as photoinitiator, and acrylated ChBT (aChBT) at concentrations of 2, 4, or 6% (w/v). Deionized water served as the solvent for all formulations unless stated otherwise. Components were added sequentially in order of decreasing water solubility to ensure complete dissolution: first, LAP was dissolved under gentle stirring, followed by ChBT, and finally GelMA was incorporated with thorough vortexing after each addition to ensure complete miscibility. The pH of the precursor solution was adjusted to 7 using 10 M NaOH to ensure physiological compatibility and ionization of the bitartrate counter-ion. The final prepolymer solution was mixed thoroughly until homogeneous and free of visible particulates. To fabricate hydrogels, 300 μL of the prepolymer solution was dispensed into the wells of a 48-well plate and cured under 405 nm irradiation for 30 s, yielding cylindrical FiBGel discs (10 mm Ø × 4 mm H). Control groups containing 0% (w/v) BIL were prepared using the same protocol but omitting the ChBT component.

### 4.4. ^1^H NMR analysis

Chemical characterization of the starting materials and hydrogels was carried out via ¹H NMR spectroscopy using a Bruker Avance 600 spectrometer operating at 600 MHz. Solutions of GelMA (7% w/v) and ChBT (4% w/v) were each prepared in deuterium oxide (D₂O, Sigma-Aldrich) and transferred to NMR tubes for direct analysis. All chemical shifts are referenced to the residual water signal of D₂O (HOD, δ 4.79 ppm). For hydrogel spectra, precursor mixtures containing GelMA (7% w/v), ChBT (4% w/v), and LAP (0.5% w/v) were dissolved in deuterium oxide under minimal ambient light, photopolymerized for 5 s using a 405 nm light source, immediately loaded into NMR tubes, and analyzed. All spectra were processed and baseline-corrected using MestReNova software (version 14.0).

### 4.5. SEM Imaging

For microstructural analysis, FiBGel samples were prepared as described above, frozen at −80 °C overnight, and then lyophilized under vacuum for 24 h to retain the native pore architecture. Cross-sectional specimens were obtained using a scalpel and mounted on aluminum specimen stubs with conductive carbon adhesive tabs. Imaging was performed using a Hitachi TM4000Plus II scanning electron microscope operating at 15 kV accelerating voltage. Initial sample positioning was conducted at low magnification for orientation, followed by high-resolution imaging for detailed morphological characterization. Images were acquired in slow scan mode utilizing BSE detection to enhance contrast and optimize visualization of the hydrogel’s microstructure.

### 4.6. EIS and Ionic Conductivity

Electrochemical impedance spectroscopy was performed on FiBGel hydrogels using a CHI660E electrochemical workstation (CH Instruments, Austin, TX) configured in a three-electrode setup (Figure 2A). The working electrode consisted of a glassy carbon electrode (GCE), with an Ag/AgCl, reference electrode and a platinum wire counter electrode. Prepolymer solutions were prepared in deionized water, dispensed into 20 mL scintillation vials to fully immerse the electrodes, and photocrosslinked in situ with 405 nm light for 30 s. Following complete polymerization, electrodes were connected to the potentiostat for impedance measurements. EIS was conducted in AC impedance mode with a 0.1 V bias, across a frequency range of 10^6^ to 10^2^ Hz, utilizing a 10 mV amplitude perturbation and a 2s quiet time. Nyquist, Bode and total impedance plots were generated. To model the electrochemical response, we fitted Nyquist data using a modified Randles circuit with constant-phase elements, in EIS Spectrum Analyzer and extracted the high-frequency solution resistance (Rs). Ohmic ionic conductivity (σ) was then obtained via cell constant (K) calibration using standard KCl solutions with known conductivity (***see SI for details***).

### 4.7. Rheological Test

The viscoelastic properties of the hydrogels were characterized using an Anton Paar rheometer equipped with a sand-blasted parallel plate geometry (25 mm diameter) to prevent wall slip. Hydrogel samples were prepared for testing by adding 2 mL of the prepolymer solution into a 25 mm diameter mold that matched the geometry, and photocrosslinking for 1 min. Once loaded onto the rheometer stage, the gap was adjusted to apply a consistent normal force of 0.1 N to ensure complete sample contact. Except for the temperature sweep experiments, all measurements were conducted at 20 °C. Frequency sweeps spanned 0.1–10 Hz at a fixed strain of 1%, a value pre-confirmed to lie within the linear viscoelastic (LVE) region. The storage modulus (G’) and loss modulus (G”) were recorded as a function of frequency. From these values, the loss tangent (tan δ), defined as the ratio G”/G’, was calculated to quantify the material’s damping behavior and overall viscoelastic character. The elastic modulus (E) was estimated from the shear modulus(G’) value at a representative frequency of 1 Hz. Assuming the hydrogel is a nearly incompressible material (Poisson’s ratio, ν ∼ 0.5), the relationship E ∼ 3G′ was used for this conversion[83]. For each formulation, a minimum of four samples were analyzed.

### 4.8. Uniaxial Compression Test

Hydrogel discs (10 mm Ø × 4 mm H) were moulded from 300 µL aliquots of pre-polymer solution in 48-well plates (Corning) and photocrosslinked as previously described. Before testing, hydrogels were allowed to reach equilibrium swelling over 4 h in PBS (pH 7.4) at 37 °C. Unconfined uniaxial compression was performed on an Instron universal testing system equipped with a calibrated 100 N load cell and polished stainless-steel platens. Each hydrogel (n ≥ 4 per formulation) was centered between the platens, a 0.001 N preload was applied to establish uniform contact, and compression proceeded at a constant cross-head speed of 1 mm min⁻¹, corresponding to an engineering strain rate of ∼ 4.2 × 10^-3^ s^-1^. Force–displacement data were recorded and converted to stress (σ) and strain (ε) using:

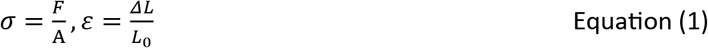

where F is the instantaneous load, A the initial cross-sectional area, ΔL the change in height, and L₀ the original height. The compressive modulus was determined as the slope of the linear region (0–10 % strain) of the resulting σ–ε curve for each sample.

### 4.9. Swelling Assay

Disc-shaped hydrogels were fabricated as outlined in the previous section, equilibrated in PBS, frozen at – 80^0^C for 24 h, and lyophilized for 24 h in a freeze dryer. The anhydrous mass of each specimen (W_dry_) was measured to the nearest 0.1 mg on an analytical balance. Lyophilized samples (n = 5 per formulation) were submerged in PBS (pH 7.4) at 37 °C and removed at 2, 4, 6, 12, and 24 h timepoints. After fluid aspiration, excess surface moisture was removed by gently wicking with lint-free tissue, and the hydrated mass (W_swollen_) was immediately recorded. The time-dependent swelling ratio was calculated as:

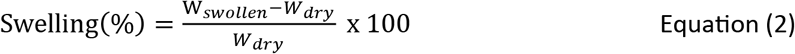

Equilibrium water content (EWC) at 24 h was calculated according to Equation 4:

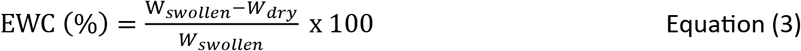

### 4.10. Hydrolytic Degradation

Hydrolytic weight loss was quantified over a 14-day interval under physiological sink conditions. Cylindrical hydrogel disks (10 mm Ø × 4 mm H) were fabricated as previously described, equilibrated in PBS, and lyophilized for 24 h in a freeze-dryer. The resulting dry weight was measured on an analytical balance and recorded as W_i_ (n = 3 per formulation per time point). Samples were then immersed in PBS (pH 7.4) and incubated at 37 °C in a humidified incubator to assess hydrolytic degradation. The medium was refreshed every 72 h to maintain sink conditions and prevent autocatalytic acidification. At predetermined time points (t = 1, 7, and 14 d), samples were gently rinsed with DI water, blotted to remove surface moisture, frozen at −80 °C, and re-lyophilized for 24 h before determining the residual dry mass W_t_. % Weight loss was calculated according to:

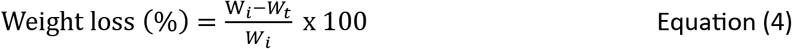

### 4.11. i-HNSC culture

Cryopreserved i-HNSCs (Cell Applications, Inc) were expanded and used between passages 4–6. Vials were thawed for 30 s in a 37 °C water bath, transferred to a 15 mL conical tube containing 9 mL prewarmed i-HNSC Growth Medium (Cell Applications), and centrifuged at 250 × g for 5 min. Cell pellets were gently resuspended in 3 mL growth medium containing 10 µM ROCK inhibitor (Y-27632 Dihydrochloride, STEMCELL Technologies) and plated at 1 × 10⁶ cells well⁻¹ in six-well plates (Corning) pre-coated sequentially with poly-L-ornithine (1 h, 37 °C) then laminin (2 h, 37 °C). After 24 h, medium was replaced with ROCK-free growth medium, and cultures were maintained at 37 °C and 5 % CO₂ in a humid incubator with complete medium exchange every third day until ∼90 % confluence. Routine passaging was performed with StemPro™ Accutase™ (Gibco) for 5 min at 37 °C and the detached cells were pelleted, resuspended in growth medium + 10 µM Y-27632, and reseeded on freshly coated plates at the same density. This workflow preserves a highly proliferative, undifferentiated i-HNSC phenotype for downstream encapsulation.

### 4.12. i-HNSC Encapsulation in FiBGel

For cytocompatibility and differentiation studies, i-HNSCs were encapsulated in engineered hydrogel matrices. 2 × precursor solutions were prepared in i-HNSC Growth Medium and sterile-filtered through 0.22 µm PES membranes (Avantor). Prior to photocrosslinking, each precursor (2 ×) was mixed 1:1 (v/v) with a concentrated cell suspension (8 × 10^6^ cells mL⁻¹), resulting in a homogeneous blend containing 4 × 10⁶ cells mL⁻¹. 30 µL aliquots were dispensed into the center of 48-well plates and polymerized via exposure to 405 nm light for 2 × 10 s with a 5 s interval to dissipate heat. Gelation was verified by gentle probe deflection. Each construct was overlaid with 400 µL i-HNSC Growth Medium supplemented with 10 µM Y-27632, and the ROCK inhibitor was removed by complete medium exchange at 1 day post-encapsulation (dpe). For differentiation assays, the growth medium was replaced with i-HNSC Differentiation Medium (Cell Applications) at 7 dpe, and cultures were maintained under standard conditions for up to 21 days post differentiation.

### 4.13. Cytocompatibility Assessments

The cytocompatibility of FiBGel hydrogels was evaluated using Live/Dead® assays, PrestoBlue® metabolic-activity measurements, and actin/DAPI staining. For Live/Dead tests, hydrogel constructs were washed twice with sterile DPBS (Dulbecco’s PBS, pH 7.4; Gibco) and then incubated in a staining solution of 2 µM calcein-AM and 4 µM ethidium homodimer-1 in DPBS. A volume of 300 µL was added per well, and the samples were incubated for 30 min in the dark at room temperature. Imaging was performed on a Zeiss LSM 880 inverted confocal microscope equipped with 488 nm (argon) and 561 nm (HeNe) lasers. Acquired Z-stacks were rendered into average-intensity 2D projections using ImageJ (NIH, Bethesda, MD, USA).

Metabolic-activity analysis (PrestoBlue®), which relies on the viability-dependent reduction of resazurin to resorufin, was conducted on days 1, 7, and 14 dpe. At each interval, the spent medium was aspirated and replaced with 300 µL of a 1:10 (v/v) dilution of PrestoBlue® in fresh i-HNSC Growth Medium. Constructs were incubated 5 h at 37 °C in a 5 % CO₂ humidified incubator. Absorbance was then recorded at 570 nm and 600 nm with a UV–vis spectrophotometer. Metabolic activity was calculated according to the manufacturer’s instructions, normalizing the A₅₇₀ values to their corresponding A₆₀₀ references and correcting against reagent-only controls (medium + PrestoBlue® without cells). All formulations were assessed in three biological replicates per time point.

To examine the morphology of i-HNSCs cultured in FiBGel hydrogels, samples were stained with fluorescently conjugated phalloidin for actin and 4′,6-diamidino-2-phenylindole (DAPI) for nucleus. Samples were washed 2x with DPBS and fixed in 4% (v/v) paraformaldehyde (PFA, Thermo Scientific) for 15 min at room temperature. After two PBS washes, cells were permeabilized with 0.1% Triton X-100 (Sigma-Aldrich) for 20 min, then non-specific binding was blocked with 1% (w/v) BSA (bovine serum albumin) for 45 min. Filamentous actin was stained using phalloidin (Cell Signaling Technology, 1:50 in 0.1 % BSA, 45 min). Nuclear counter-staining was performed with DAPI (1:1000 in PBS, 5 min), followed by three PBS washes. Z-stack confocal images were acquired, processed into 2D projections, and morphometric analysis was conducted using ImageJ to determine cell length (µm).

### 4.14. Immunostaining

Cell-laden hydrogels were processed to visualize early (β3-tubulin) and mature (MAP2) neuronal markers. Constructs were rinsed twice in sterile 1× PBS and fixed for 15 min at room temperature in 4% (v/v) PFA (Thermo Scientific). After three washes in ice-cold PBS, samples were permeabilized in 0.3 % Triton X-100 (Sigma-Aldrich) for 15 min, followed by two further PBS washes (5 min each). Non-specific binding sites were blocked in 1% (w/v) bovine serum albumin (BSA) + 0.1 % Triton X-100 for 1 h. Primary antibodies were prepared in blocking buffer at the concentrations specified. For neurogenic assays: rabbit anti–MAP2 (1:400; Cell Signaling Technology, #4542) and mouse anti–βIII-tubulin (1:200; Invitrogen, MA1-118). For angiogenesis assays: mouse anti–CD31 (1:300; Abcam, ab24590), rabbit anti–VE-cadherin (1:300; Abcam, ab33168), and rabbit anti–vWF (1:300; Abcam, ab6994). Hydrogels were incubated overnight at 4 °C in a humidified chamber, then washed three times with PBS (5min each). Secondary labelling was carried out with goat anti-mouse IgG Alexa Fluor 594 (1:400, Invitrogen, A-11032) and goat anti-rabbit IgG Alexa Fluor 488 (1:400, Invitrogen, A-11034) for 2 h at room temperature, protected from light. Excess secondary antibody was removed via three PBS washes. Nuclei were counter-stained with DAPI (1:1000 in PBS, 5 min) followed by a final PBS rinse. Fluorescence images were acquired on a Zeiss LSM 880 inverted confocal microscope, and imaging parameters were kept consistent for all samples.

### 4.15. Calcium Imaging

Intracellular calcium dynamics were recorded on day 21 post-differentiation using the acetoxymethyl ester probe Fluo-8® AM (AAT Bioquest). A 5 mM stock was prepared by dissolving 50 µg lyophilized dye in 10 µL anhydrous dimethyl-sulfoxide (Sigma-Aldrich). Before use, the stock was diluted in i-HNSC differentiation medium to 5 µM Fluo-8® AM and supplemented with 0.04 % (v/v) Pluronic® F-127 to enhance dispersion. Spent medium was aspirated and replaced with 300 µL dye solution. Afterwards, constructs were maintained for 30 min at 37 °C, 5% CO₂ to permit uptake and de-esterification, then rinsed once with Hanks’ Hepes-buffered saline (HHBS; Gibco). Time-lapse imaging was performed on the Zeiss LSM 880 confocal microscope equipped with an FITC filter set (excitation = 490 nm, emission = 525 nm, cutoff = 515 nm). Time-lapse recordings were captured for 120 cycles at 1.26 s intervals (total duration ∼ 150 s). Image frames were exported in CZI, TIFF, and MOV formats. ROIs corresponding to spontaneously active neurons were tracked in ImageJ, and fluorescence traces normalized considering F_t_/F_0_, where F_0_ is the baseline fluorescence averaged over quiescent frames and F_t_ is the fluorescence at time t.

### 4.16. DLP Bioprinting of Cell-Laden FiBGel

A CELLINK Lumen X Gen 1 DLP bioprinter (405 nm) was employed to fabricate both acellular and cell-laden constructs. Acellular bioinks were formulated in PBS containing 7% (w/v) GelMA, 0.25% (w/v) LAP, 4% (w/v) choline bitartrate, and 0.02% (w/v) tartrazine as a photo-absorber. Printing was performed on hydrophobic PDMS-coated build plates using optimized parameters: 60% power (∼18.8 mW/cm²), 8 s layer exposure, 1× burn-in factor, and 50 μm layer height. Post-printing, constructs were rinsed in DI water for 30 s to remove uncured resin. For cell-encapsulated constructs, PBS was replaced with i-HNSC growth medium and tartrazine was omitted to mitigate cytotoxicity. i-HNSCs were mixed into the sterile filtered bioink at a final density of 4 × 10⁶ cells/mL. Printing proceeded under sterile conditions using parameters adjusted for the medium’s intrinsic attenuation: 35% power (∼10.96 mW/cm²), 3 s exposure, 1× burn-in factor, and 100 μm layer height. Printed constructs were detached using a sterile blade and transferred to 48-well plates with 400 μL of growth medium. All printed constructs were visually inspected to confirm the absence of delamination and over-polymerization before downstream assays.

### 4.17. HUVEC culture

HUVECs were expanded in Endothelial Growth Medium-2 complete medium (EGM-2; basal medium plus BulletKit supplements) (CC-3162, Lonza, Walkersville, MD). Cryovials were thawed for 30 s in a 37 °C water bath, transferred to 5 mL of pre-warmed EGM-2 in a 15 mL conical tube and centrifuged at 350 × g for 5 min. The cell pellet was gently resuspended in fresh EGM-2 and plated at 8 × 10⁵ cells per T-75 flask (Thermo Scientific). Medium was exchanged every other day until cultures reached ∼80 % confluence (3– 4 d). For routine passaging, monolayers were rinsed with PBS, detached with 0.25% (w/v) trypsin–EDTA (Gibco) for 2–3 min, neutralized with EGM-2, and centrifuged at 300 × g for 5 min before reseeding at a 1:4 split ratio. For cryopreservation, detached cells were suspended in EGM-2 supplemented with 10% (v/v) fetal bovine serum and 10% (v/v) DMSO, cooled at −1 °C min⁻¹ to −80 °C, and subsequently stored in liquid nitrogen. This workflow consistently maintained a highly proliferative, undifferentiated HUVEC phenotype suitable for subsequent 3D encapsulation studies.

### 4.18. In Vitro Angiogenesis Assay in FiBGel Matrices

HUVECs at passage 6 were encapsulated in FiBGel hydrogels following the same workflow used for i-HNSC experiments. Briefly, 2 × precursor solutions were prepared in EGM-2 medium and sterilized through 0.22 µm PES filters. Immediately before photocrosslinking, each precursor was mixed 1:1 (v/v) with a concentrated HUVEC suspension (4 × 10⁶ cells mL⁻¹), yielding a homogeneous 2 × 10⁶ cells mL⁻¹ solution. Aliquots of 30 µL were deposited in the center of non-treated 48-well plates and cured by two 10 s pulses of 405 nm light with a 5 s inter-pulse interval to prevent heat accumulation. Gelation was confirmed by gentle probe deflection. Each construct was then overlaid with 400 µL EGM-2 and maintained at 37 °C/5% CO₂ for 21 days, with medium replenished every 48 h. Endothelial network formation was monitored throughout the culture by phase-contrast microscopy and assessed at the study endpoint by immunostaining for CD31, VE-cadherin, and vWF.

### 4.19. Lateral Fluid Percussion Injury and Hydrogel Administration

All animal procedures were conducted in compliance with protocols approved by the Institutional Animal Care and Use Committee (IACUC) of the University of California, Riverside, and following the ARRIVE guidelines for reporting animal research. Male and female C57BL/6J littermate mice (8–10 weeks old, n = 3 per group) were randomly assigned to receive either lateral fluid percussion injury (FPI) or sham surgery[84]. Mice were anesthetized with isoflurane and placed in a stereotaxic apparatus. Local anesthesia was provided with 0.25% bupivacaine administered subcutaneously at the incision site. A 2.7 mm craniectomy was drilled over the left hemisphere, centered 2 mm posterior to bregma and 1 mm lateral to the sagittal suture. A Luer Lock syringe hub (inner diameter 3 mm) was affixed to the exposed dura using cyanoacrylate adhesive. After recovery from anesthesia, mice were returned to their home cages. 24 h later, mice were re-anesthetized and connected to a fluid percussion device (Model 01-B, Virginia Commonwealth University, Richmond, VA). A pendulum was released to deliver a rapid (20 ms) fluid pressure pulse of 1.5–1.7 atm directly to the intact dura via the hub. Sham-operated mice underwent identical procedures, including hub implantation and connection to the FPI device, but without pendulum release[60,61,84].

Immediately following FPI or sham-injury, mice received stereotaxic hydrogel injections into the lesion site under anesthesia. Treatments included FiBGel, GelMA, or saline as vehicle controls. Using a Hamilton syringe coupled to a microinjector, 100 µL of hydrogel or vehicle solution was injected at the following stereotaxic coordinates relative to bregma: anteroposterior (AP) −2.0 mm, mediolateral (ML) −1.0 mm, dorsoventral (DV) −0.5 mm. Injections were performed slowly over 5 minutes to minimize reflux. For treatments, *in situ* crosslinking was achieved using a 405 nm blue light directed at the injection site for ∼1 min. Vehicle controls were also exposed to 405 nm blue light directed at the injection site for ∼1 min. Following injections, the hub implant was removed, the scalp was closed with tissue adhesive, and mice were monitored until recovery from anesthesia before being returned to their home cages.

### 4.20. Tissue Processing and Immunohistochemistry

At 7 days post-injury (dpi), mice were transcardially perfused first with ice-cold PBS and then with 4% PFA in PBS. Brains were removed, post-fixed in 4% PFA overnight at 4 °C, and cryoprotected in 30% sucrose until the tissue sank (2–3 days)[84]. Samples were then embedded in optimal cutting temperature (OCT) compound, rapidly frozen using liquid nitrogen, and stored at −80°C until cryosectioning. Serial coronal sections (20 μm thickness) were obtained using a cryostat, mounted onto SuperFrost™ slides, and stored at −80°C until further use.

For immunostaining, sections were equilibrated to room temperature for 10 minutes and washed 3 times (5 minutes each) in PBS to remove residual OCT. Sections were permeabilized with 0.5% Triton X-100 for 45 min, then blocked in 20% normal goat serum (Invitrogen) containing 0.025% Triton X-100 in PBS for 1 h at room temperature, followed by incubation with primary antibodies. These included mouse anti-Iba1 (1:300, Sigma-Aldrich, MABN92), rabbit anti-GFAP (1:200, Cell Signaling, 80788), or mouse anti-CD31 (1:300, Abcam, ab24590) overnight at 4^0^C. The following day, slides were washed in PBS (3 × 10 minutes) and incubated with appropriate fluorescent secondary antibodies: goat anti-mouse IgG Alexa Fluor 594 (1:400, Invitrogen, A-11032) or goat anti-rabbit IgG Alexa Fluor 488 (1:400, Invitrogen, A-11034) for 2 h at room temperature. After 3 PBS washes (10 minutes each), sections were coverslipped using VectaShield® antifade mounting medium with DAPI.

Images were acquired using a Nikon fluorescent microscope. Cortical and hippocampal regions from both hemispheres were captured via stitched 6×2 tile acquisition using standardized field overlap and blending parameters to ensure seamless montage reconstruction. Higher magnification images were obtained at 20× using a Zeiss LSM 880 inverted confocal microscope. Imaging parameters were kept constant across all experimental conditions to ensure valid quantitative comparisons. Regional fluorescence intensity (mean gray value) measurements were performed using ImageJ software with ROI tracings for target anatomical areas.

### 4.21. Statistical Analysis

Statistical analyses and graphical representations were performed using GraphPad Prism version 10.0 (GraphPad Software, Boston, MA, USA). Experimental data are expressed as mean ± standard error of the mean (SEM) from a minimum of three independent biological replicates, unless otherwise stated. Comparative analyses between two experimental groups were conducted using unpaired two-tailed Student’s t-tests. For multiple group comparisons involving a single independent variable, one-way analysis of variance (ANOVA) was employed, followed by Tukey’s post hoc multiple comparisons test to identify specific pairwise differences. When analyzing datasets with two independent variables, two-way ANOVA was utilized with subsequent Tukey’s multiple comparisons testing. Statistical significance thresholds were established at *p < 0.05, **p < 0.01, ***p < 0.001, and ****p < 0.0001.

## Data Availability Statement

The authors declare that all data supporting the findings of this study are available within the manuscript and its Supplementary Information. Additional data related to this work are available from the corresponding author upon reasonable request.

## Author Contributions

P.D.O. contributed to conceptualization, data curation, formal analysis, funding acquisition, investigation, methodology, software development, validation, visualization, and writing the original draft, review, and editing. S.B. contributed to methodology and investigation. R.J. contributed to methodology, investigation, and writing, review, and editing. D.P. contributed to methodology and investigation. J.M., M.S., R.B., and J.T. contributed to methodology. V.I.V. and H.L. contributed to writing, review, and editing. V.S. contributed to investigation, writing, review, and editing. I.N. contributed to conceptualization, funding acquisition, investigation, methodology, project administration, resources, supervision, validation, writing, review, and editing.

## Acknowledgements

The authors gratefully acknowledge financial support from the UCR startup fund. P.D.O. is supported by a TRANSCEND fellowship from the California Institute of Regenerative Medicine (Award # EDUC4-12752). The contents of this publication are solely the responsibility of the authors and do not necessarily represent official views of CIRM or other agencies of the State of California.

