## Supplementary Information for "A Neurocompliant Ionic Conductive Hydrogel Orchestrates Glial and Vascular Remodeling to Mitigate Traumatic Brain Injury"

R. J., Prof. V. S.
Department of Molecular, Cell and Systems Biology, University of California Riverside, Riverside, CA, USA

Keywords: Bioionic liquid, Brain tissue engineering, Neuroinductive materials, Angiogenesis, Astroglial modulation, Traumatic brain injury

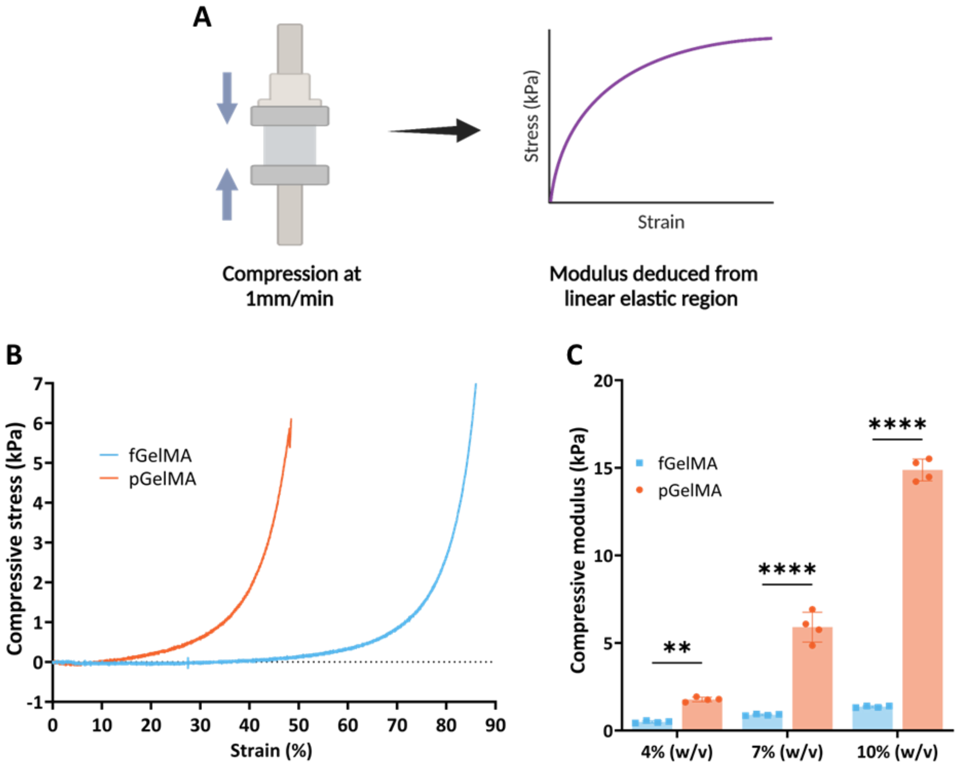


Figure S1. **Mechanical properties of fish and porcine GelMA hydrogels.** (A) Schema of unconfined compression test: cylindrical hydrogel disks were compressed at 1 mm $min^{-1}$, and the compressive modulus was calculated from the linear elastic region of the stress–strain curve. (B) Representative compressive stress–strain curves for 4 wt% fish gelatin methacryloyl (GelMA) and porcine GelMA, highlighting the stiffer response of porcine GelMA. (C) Compressive modulus of fish GelMA and porcine GelMA at 4, 7, and 10% (w/v); porcine GelMA exhibited significantly higher moduli at equivalent concentrations. Values are mean ± SEM. Statistical significance via two-way ANOVA with Tukey's post-hoc test (ns: not significant, ***p* ≤ 0.01, *****p* ≤ 0.0001).

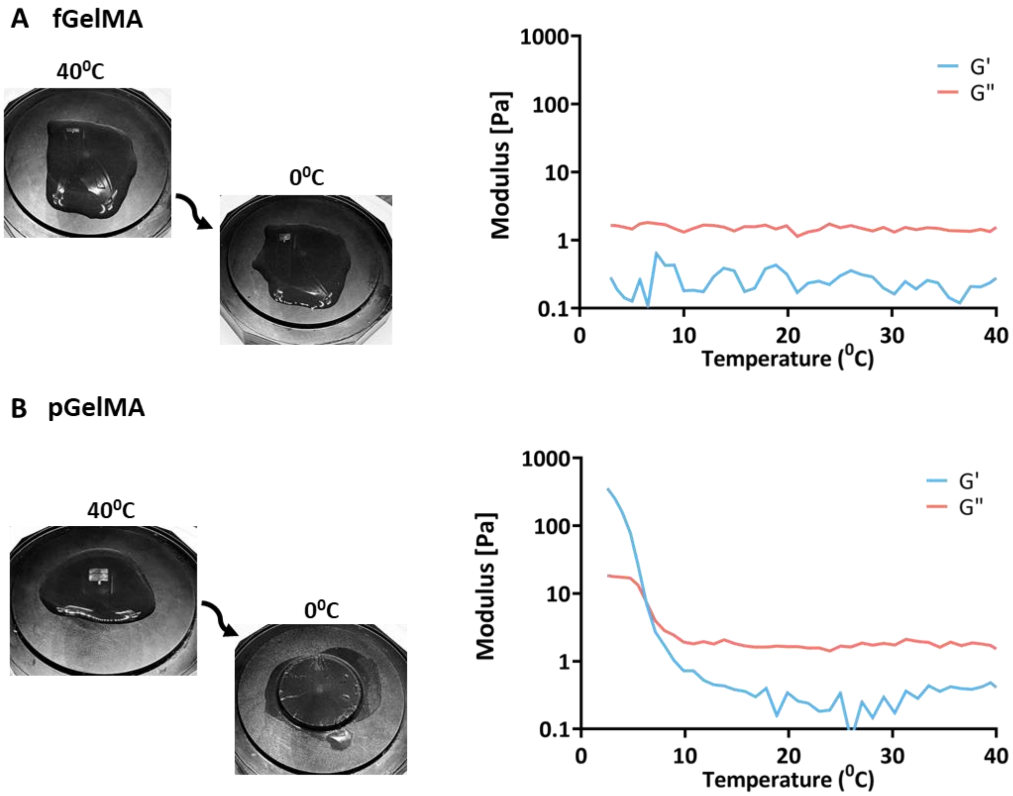


Figure S2. **Thermoresponsive gelation of fish and porcine GelMA.** (A) Representative images and rheological analysis of fish gelatin methacryloyl (GelMA) solution, showing G′ (shear modulus) and G″ (loss modulus ) across a temperature range of 40°C to 0°C. G′ remains relatively stable, indicating minimal thermal sensitivity. (B) Images and temperature-dependent rheological analysis of porcine GelMA solution, with G′ increasing sharply above 10°C, highlighting thermoresponsive sol-gel transition. Both moduli (G′ and G″) are plotted on a logarithmic scale.

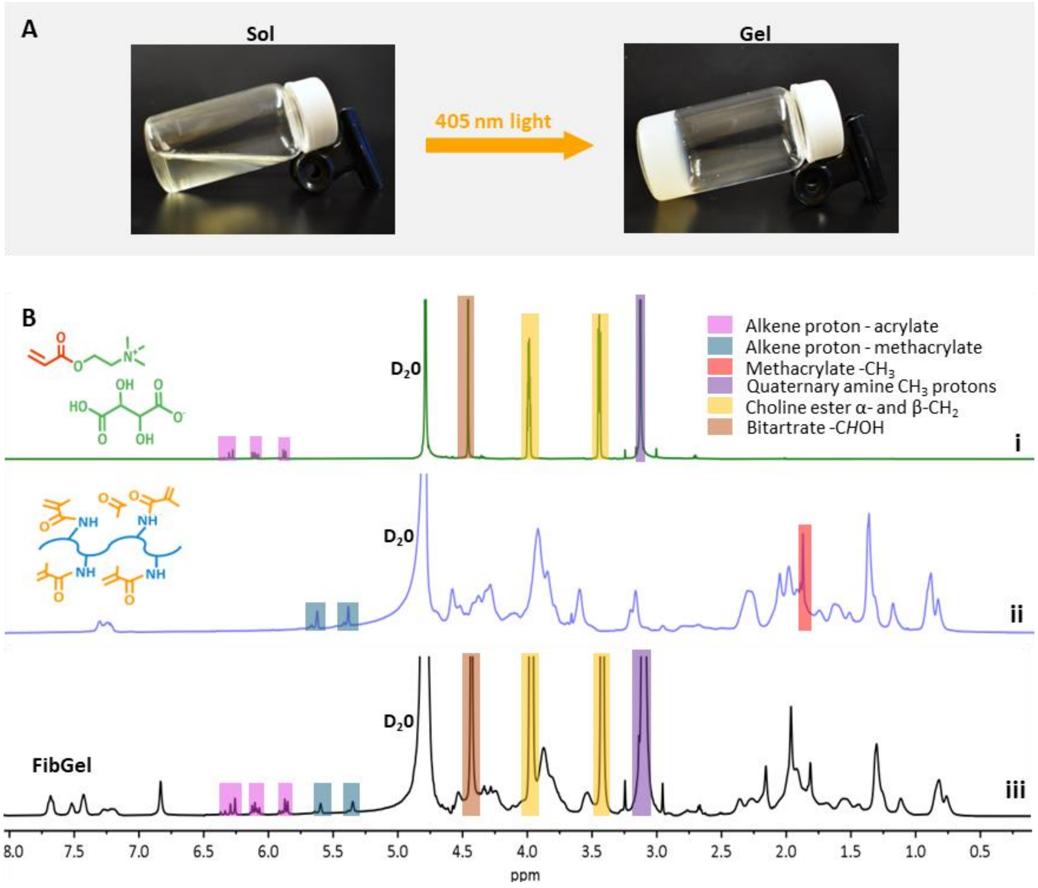


Figure S3. **Gelation and chemical characterization of FiBGel hydrogels**. (A) Visual representation of the sol–gel transition after 405 nm irradiation for 30 s, shown for bioionic liquid (BIL)-conjugated fish gelatin methacryloyl (GelMA) hydrogels (FiBGel). (B) $^{1}$H NMR spectra of (i) acrylated choline bitartrate (ChBT), (ii) fish GelMA, and (iii) FiBGel, recorded at 600 MHz (Bruker Avance 600) in deuterium oxide ($D_2O$). Chemical shifts (δ) are reported in ppm relative to the residual solvent peak (HOD, δ 4.79 ppm).

**Supplementary Note 1: Electrochemical impedance spectroscopy**

To estimate the ionic conductivity of hydrogels, we resort to electrochemical impedance spectroscopy (EIS). Applying a small AC signal across a wide frequency range provides a means for deconvoluting the response from the hydrogel into bulk and interfacial contributions. It allows us to extract the bulk ohmic solution resistance ($R_S$) and compute ionic conductivity $\sigma$. We tested hydrogels in a three-electrode cell with a glassy-carbon working electrode, an Ag/AgCl reference, and a platinum counter electrode (Figure S4A), using a 10 mV perturbation in the range from about $10^6$ to $10^2$ Hz. Nyquist plots display the real part of impedance (Z′) on the x-axis and the negative imaginary (−Z″) on the y-axis (Figure S4B). The high-frequency left intercept on the Nyquist plot corresponds to Rs, while a depressed arc at mid frequencies indicates interfacial polarization. Bode magnitude and phase plots complement the Nyquist representation of the data by showing how impedance and phase evolve with frequency. These assignments follow standard EIS practice[1,2] and give a robust readout of bulk ion transport and interface behavior.

To extract $R_S$, Nyquist plots were fitted using a modified Randles circuit with constant phase elements, R(Q(RQ)) as shown in Figure 1D. This model reproduces the depressed semicircle and the sub-diffusive tail without biasing $R_S$. In this model, $R_S$ sits in series with a parallel network where the upper arm is a CPE that captures the nonideal double layer, and the lower arm contains the charge-transfer resistance, $R_{ct}$, in series with a second constant-phase element (CPE). Fittings were performed in EIS Spectrum Analyzer, which uses complex nonlinear least-squares algorithm with amplitude weighting. Only $R_S$ was used for computing $\sigma$. We converted extracted $R_S$ values to conductivity by using a cell constant ($K$) obtained from calibration with KCl standards of known conductivity. Specifically, we measured 0.01 M KCl at 25 ± 1 °C, read $R_S$ for KCl, and set $K = R_{s,\mathrm{KCl}}\sigma_{\mathrm{KCl}}$ considering the certified conductivity of 0.1413 S $m^{-1}$[3,4]. We then convert each hydrogel $R_{s,\mathrm{gel}}$ (Figure S4C) to ionic conductivity using $\sigma_{\mathrm{gel}} = K/R_{s,\mathrm{gel}}$. This approach removes uncertainty in electrode spacing and wetted area and ties the measurement to traceable NIST standards[3,4]. Linearity was verified by repeating the calibration with 0.1 M and 1.0 M KCl.

A

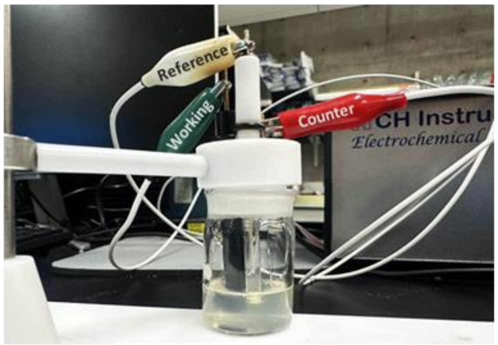


B

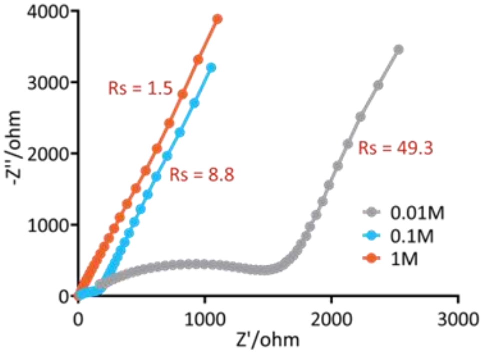


C

| | 2% BT | 4% BT | 6% BT | GelMA |
|---|---|---|---|---|
| $C_{BIL}$ / % (w/v) | 2 | 4 | 6 | 0 |
| *Rs* / Ω | **27.88±1.69** | **12.95±1.15** | **10.30±1.21** | **169.58±10.48** |

Figure S4. **EIS analysis of FiBGel hydrogels.** (A) A three-electrode electrochemical setup employed for EIS, comprising: (1) a glassy-carbon working electrode, (2) an Ag/AgCl reference electrode, and (3) a platinum counter electrode. (B) Nyquist plots of KCl standards at 25 °C. The high frequency intercept ($R_S$) defines the cell constant used for converting the ohmic resistance of the hydrogel to conductivity. (C) Extracted *Rs* (mean ± SD) for FiBGel formulated with choline bitartrate (ChBT) at 2, 4, or 6% (w/v) and for GelMA control, showing a concentration-dependent drop in $R_S$.

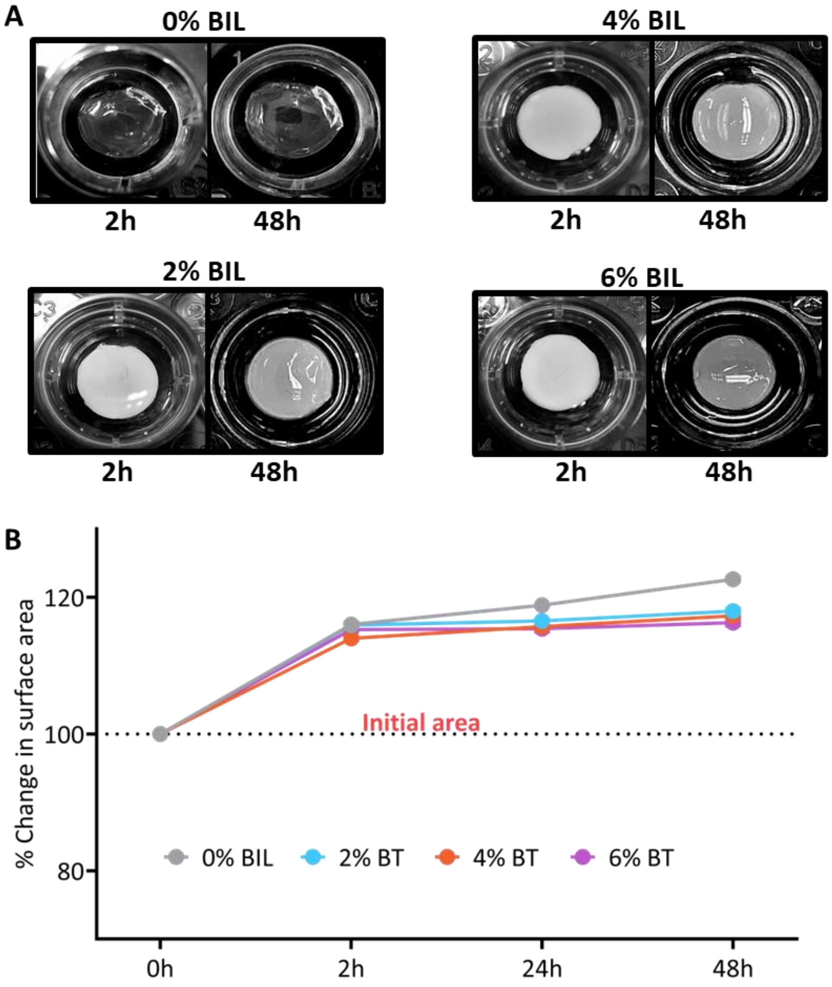


Figure S5. **Stability assessment of FiBGel hydrogels.** (A) Representative photographs of bioionic liquid (BIL)-conjugated fish gelatin methacryloyl (GelMA) hydrogels (FiBGel) functionalized with choline bitartrate (ChBT) (0–6% w/v) at 2 h and 48 h. (B) Percentage change in surface area of FiBGel hydrogels over 0, 2, 24, and 48 h, demonstrating structural stability upon BIL incorporation. Data are mean ± SEM.

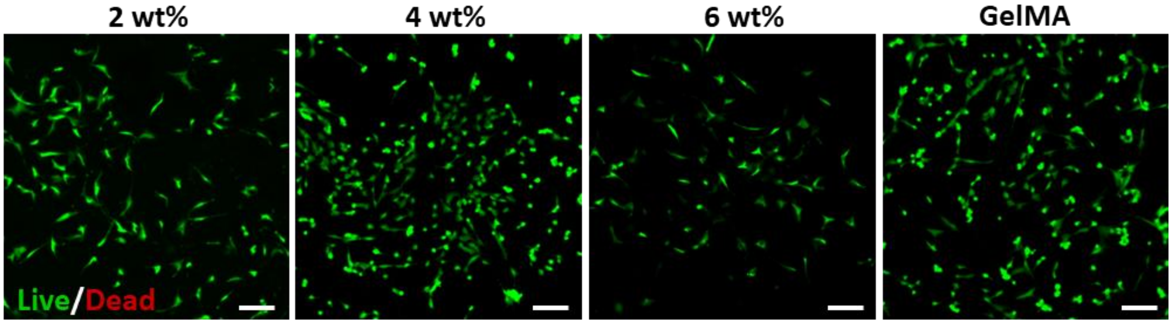


Figure S6. **Viability of i-HNSCs in FiBGel hydrogels on Day 7.** Live/Dead staining of iPSC-derived human neural stem cells (i-HNSCs) encapsulated in bioionic liquid (BIL)-conjugated fish gelatin methacryloyl (GelMA) hydrogels (FiBGel). Formulations contained choline bitartrate (ChBT) at 2, 4, or 6% (w/v) alongside a GelMA control. Live cells are stained with calcein-AM (green), and dead cells are stained with ethidium homodimer-1 (red). Scale bars: 100 µm.

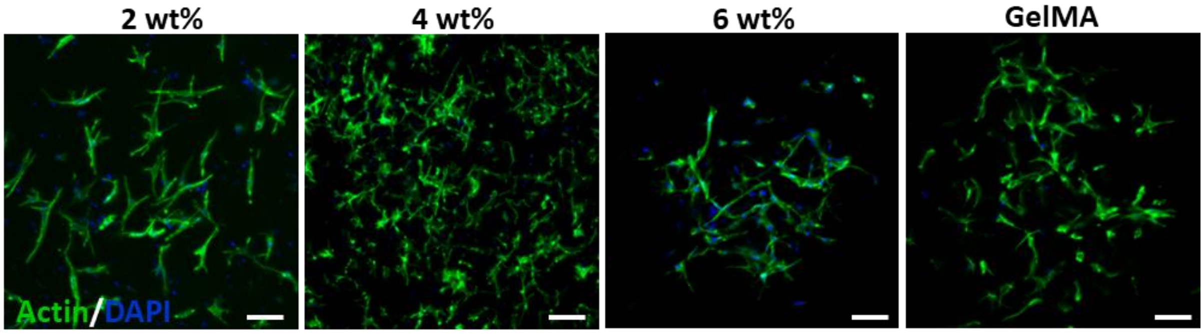


Figure S7. **Morphological assessment of i-HNSCs in FiBGel hydrogels on Day 7.** Confocal micrographs showing F-actin (green) and nuclei (DAPI, blue) staining of iPSC-derived human neural stem cells (i-HNSCs) encapsulated in bioionic liquid (BIL)-conjugated fish gelatin methacryloyl (GelMA) hydrogels (FiBGel). Formulations consisting of choline bitartrate (ChBT) at 2, 4, and 6% (w/v) were compared against a GelMA control. Scale bars: 100 µm.

**Supplementary Note 2: Photopolymerization Kinetics and Bioprinting Optimization**

Successful stereolithographic printing of hydrogels requires precise control over photopolymerization kinetics, specifically the balance between cure depth ($C_d$) and light scattering. According to the Beer-Lambert law for vat polymerization[5,6], $C_d$ is governed by the energy dose ($E$) relative to the critical energy ($E_c$) and penetration depth ($D_p$): $C_d = D_p \ln(E/E_c)$

For the acellular FiBGel formulation, the hydrogel precursors (GelMA and Choline Bitartrate) are largely transparent to 405 nm light, leading to excessive $C_d$ and uncontrolled lateral over-polymerization (light bleeding). To resolve this, we introduced tartrazine as a biocompatible photo-absorber to reduce $D_p$ and confine crosslinking to the targeted voxel height. We performed systematic parameter sweeps using lattice calibration grids (Figure S8) to identify the optimal window where structural fidelity was maximized without delamination. Power levels were varied from 50% to 80% and exposure times from 7 to 10 s. We determined that a concentration of 0.02% (w/v) tartrazine with an energy dose of ~18 $mW/cm^2$ (60% power for 8 s) yielded the sharpest features for the 50 µm layer height.

For cell-encapsulated constructs, tartrazine was omitted to eliminate potential risks of small-molecule cytotoxicity. In its place, we leveraged the intrinsic molar extinction coefficients of the phenol red and proteins present in the i-HNSC growth medium to act as photo-attenuators. However, as the absorption efficiency of the medium differs from that of the tartrazine-doped saline, the synthesis parameters required recalibration. We observed that maintaining the high energy used for acellular prints compromised cell viability, while insufficient energy led to layer delamination during the Z-lift of the build platform. Optimization revealed that increasing the layer height to 100 µm while reducing the irradiance to ~11 $mW/cm^2$ (35% power) and exposure time to 3 s provided an ideal balance. This relaxed protocol results in high viability while maintaining sufficient gelation strength to support the 3D architecture (Figure S9). To further mitigate mechanical stress during the lift-release cycle, build plates were coated with hydrophobic PDMS, reducing the adhesion force at the vat interface.

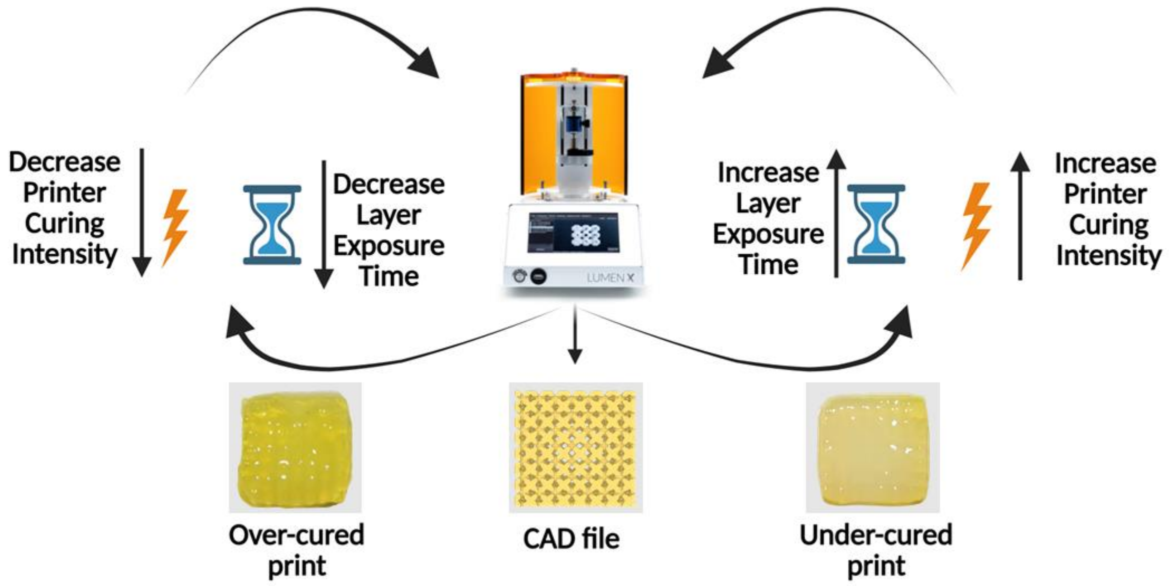


Figure S8. **Optimization of FiBGel as a bioink for DLP bioprinting.** Schematic illustrating the optimization of printing parameters for digital light processing (DLP). To achieve structural fidelity close to the CAD file, iterative adjustments are made to printing parameters. Over-curing is mitigated by decreasing light intensity or exposure time, while under-curing is corrected by increasing these same parameters. Tested bioink composition: 7% (w/v) fish gelatin methacryloyl (GelMA), 0.25% (w/v) lithium phenyl-2,4,6-trimethylbenzoylphosphinate (LAP), 4% (w/v) choline bitartrate, and 0.02% (w/v) tartrazine as a photoabsorber.

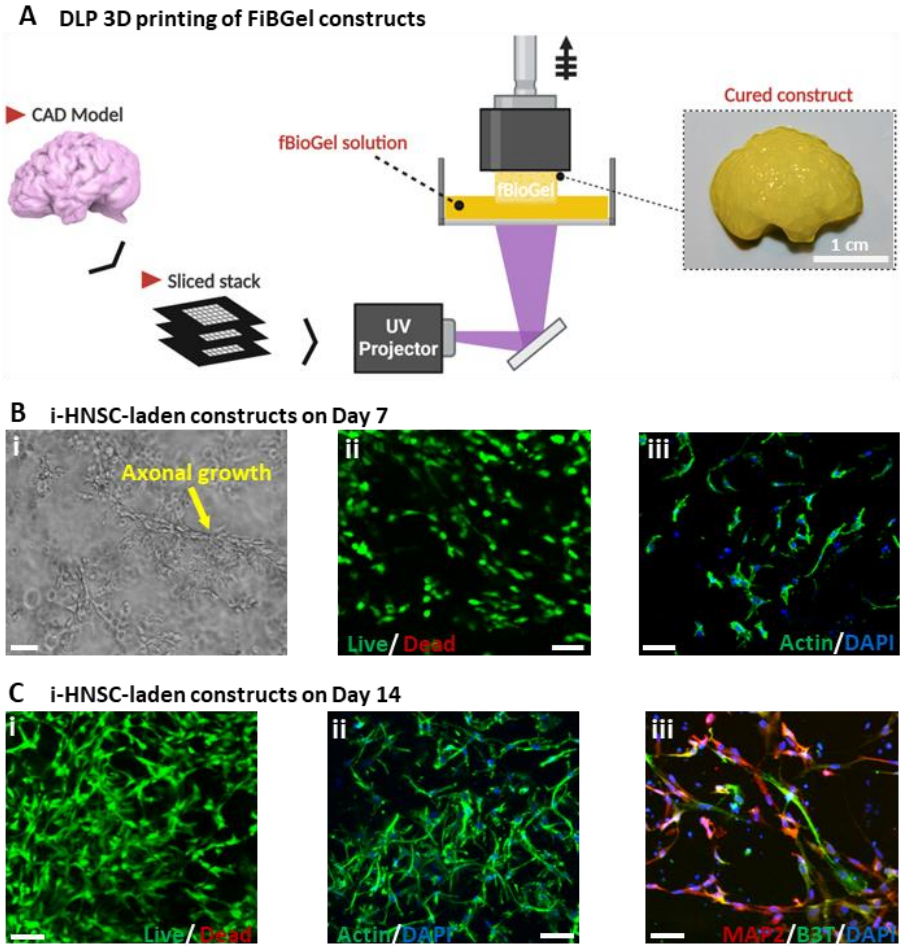


Figure S9. **Bioprinting cell-laden FiBGel constructs**. (A) Schema of the digital light processing (DLP) printing workflow includes a computer-aided design (CAD) model is sliced into layers, where each layer is selectively photocured with 405 nm light to produce a centimetre-scale, bioionic liquid-conjugated fish gelatin methacryloyl (GelMA) hydrogel (FiBGel) construct. Bioink composition: 7% (w/v) GelMA, 4% (w/v) choline bitartrate (ChBT), 0.25% (w/v) lithium phenyl-2,4,6-trimethylbenzoylphosphinate (LAP), 0.02% (w/v) tartrazine. (B) Constructs encapsulating iPSC–derived human neural stem cells (i-HNSCs) after 7 days of culture. (i) Phase contrast image showing axonal extensions (yellow arrow). (ii) Live/Dead staining (calcein AM, green; ethidium homodimer-1, red) confirms high viability. (iii) Phalloidin labelled F-actin (green) with DAPI-stained nuclei (blue) reveals early cytoskeletal organization. Tartrazine was replaced with culture media for biological studies. (C) i-HNSC-laden constructs after 14 days. (i) Live/Dead staining demonstrates sustained viability and increased cell density. (ii) Actin/DAPI images illustrate extensive neurite networks. (iii) Immunostaining for microtubule associated protein-2 (MAP2, red) and β3-tubulin (β3T, green) confirms neuronal differentiation and maturation.

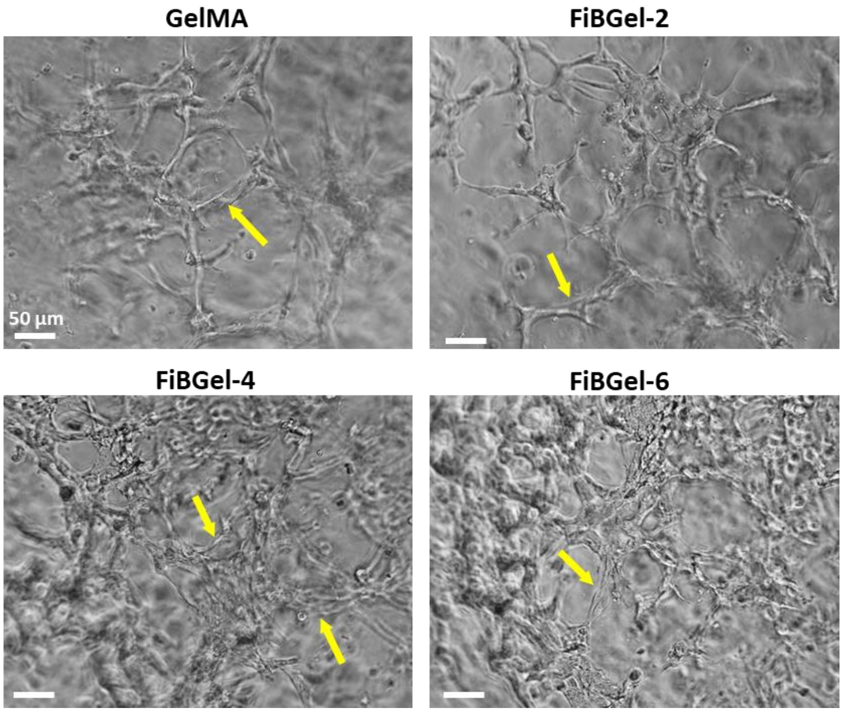


Figure S10. **Capillary-like network formation in FiBGel hydrogels on Day 14.** Phase-contrast micrographs show human umbilical vein endothelial cells (HUVECs) forming networks within fish gelatin methacryloyl (GelMA) control hydrogels and in bioionic liquid (BIL)-conjugated GelMA (FiBGel) containing 2, 4, or 6% (w/v) choline bitartrate (ChBT) (labelled FiBGel-2, FiBGel-4, and FiBGel-6, respectively). Yellow arrows highlight capillary-like tubular structures, which become denser and more interconnected as ChBT concentration increases. Scale bars: 50 µm.

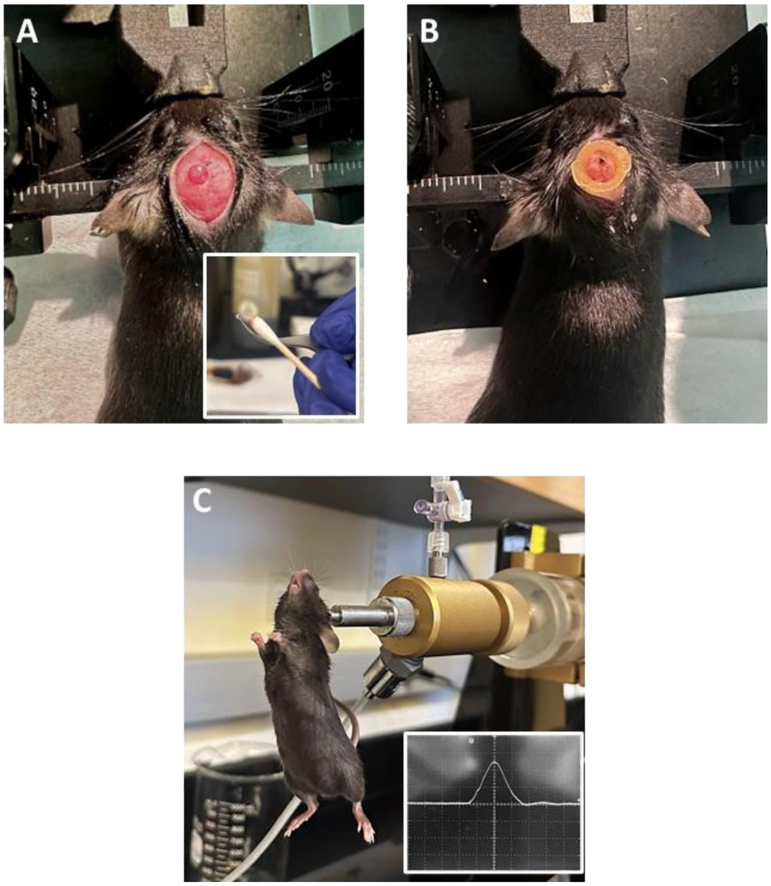


Figure S11. **Traumatic brain injury induction using a fluid percussion injury model.** (A) Following securing the skull in a stereotaxic frame, a 3 mm parietal craniectomy is made leaving the dura intact (inset: excised bone flap). (B) A Luer Lock hub is cemented over the craniotomy to create a sealed fluid column for impact delivery. (C) The hub–animal assembly is coupled to the fluid percussion injury (FPI) device, aligned beneath the piston, and subjected to a calibrated pressure pulse; the inset displays characteristic FPI waveform with a smooth rise.

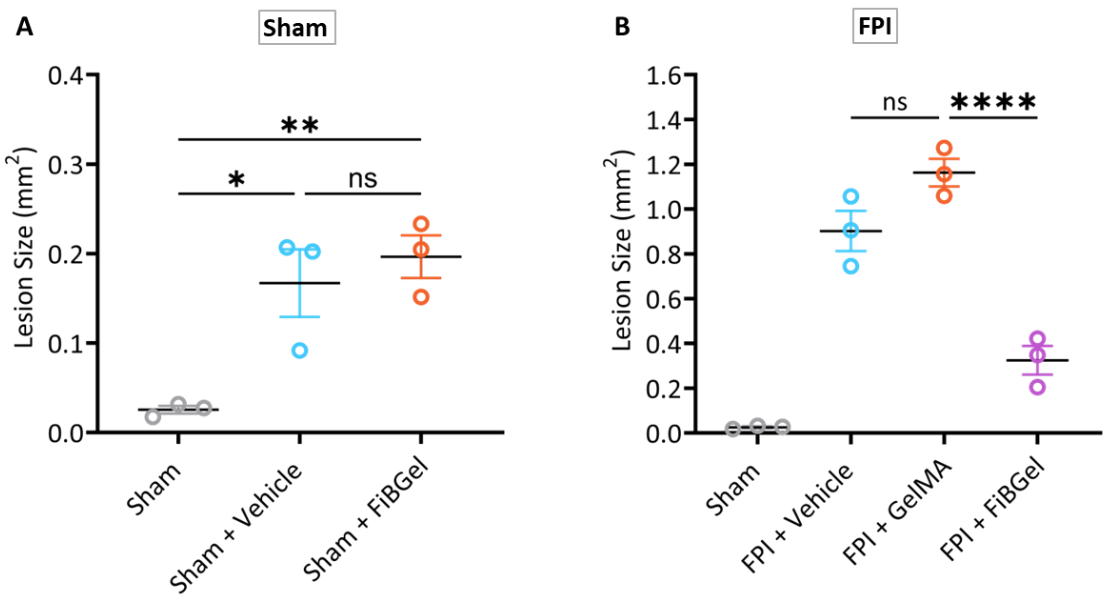


Figure S12. **Lesion cavity area in sham and FPI groups.** (A) Lesion cavity areas quantified from DAPI-stained coronal brain sections 7 days post-injury (dpi) in sham groups. (B) Corresponding cavity areas in mice subjected to moderate fluid percussion injury (FPI). Areas were manually traced in ImageJ, averaged per animal, and plotted as individual data points with group means ± SEM (n = 3). Statistical significance was determined by one-way ANOVA with Tukey's post-hoc test (ns: not significant, *$p \leq 0.05$, **$p \leq 0.01$, ****$p \leq 0.0001$).

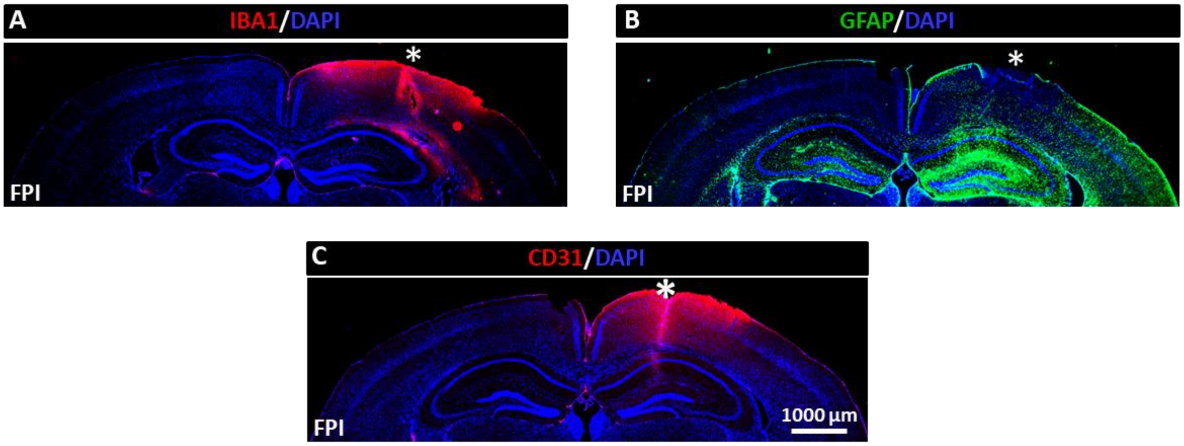


Figure S13. **Immunohistochemical analysis of cellular responses following moderate fluid percussion injury.** Representative coronal brain sections at 7 days post-injury showing distinct pathological responses at the impact site (asterisk). (A) Iba1 immunostaining (red) reveals activated microglia concentrated around the injury epicenter. (B) GFAP immunofluorescence (green) demonstrates reactive astrogliosis in different regions. (C) CD31 staining (red) highlights endothelial cells and microvascular disruption in peri-lesional regions. Nuclear counterstaining with DAPI (blue) provides anatomical reference. CD31, cluster of differentiation 31; GFAP, glial fibrillary acidic protein; Iba1, ionized calcium-binding adapter molecule 1.
